# Spatial drug asymmetry modulates phenotypic diversity-migration relationships under resistance evolution

**DOI:** 10.1101/2024.12.16.628583

**Authors:** Zhijian Hu, Yuzhen Wu, Kevin Wood

**Author notes:** Deceased (Kevin Wood).

## Abstract

At long timescales, resistant phenotypes will emerge and be selected within the bacterial population as an evolutionary response to drug exposure. This phenomenon reduces the efficacy of drug therapies and thus compromises patient health. In spatially heterogeneous drug environments, recent evidence shows that migration can either promote or decelerate the evolution of antibiotic resistance, thereby affecting the rate of resistant phenotype emergence. However, another important quantitative aspect of resistance evolution—bacterial phenotypic diversity—has often been overlooked and remains challenging to investigate in spatially extended systems, both experimentally and clinically. In order to study how diversity is reshaped by migration across space, here we designed a minimal 2-well experimental system with spatial drug asymmetry. One well contained a bacteriostatic drug (Linezolid) at the minimum inhibitory concentration, while the other well served as a sanctuary with just media. We found that the relationship between diversity and migration follows the “Intermediate Disturbance Hypothesis” (IDH), with migration as the disturbance to each well. By varying the selective drug concentrations, we observed that the diversity-migration relationship changes, and IDH can disappear. This behavior was explained by an asymmetry parameter derived from a two-phenotype growth-migration dynamic model.

To further validate how different spatial drug asymmetries modulate the diversity-migration relationship through this asymmetry parameter, we applied another bactericidal drug, Ampicillin, and observed similar results. In a more complex scenario involving both Linezolid and Ampicillin, four distinct phenotypes, including cross-resistant variants, emerged. Our asymmetry parameter successfully explained the diversity-migration relationship, with unique diversity dynamics such as multiple peaks appearing from the model. The minimal generalist-specialist framework predicted these unique behaviors through global fitness advantages. Our findings provide experimental support and theoretical explanations for the emergence of phenotypic diversity in clinical settings, such as gut-lung translocation. These insights may pave the way for improved clinical strategies to manage antibiotic resistance evolution.

## Introduction

Drug resistance has become a significant and rapidly growing public health threat, posing a major challenge to the effective treatment of cancer, viral infections, and microbial diseases [1–4]. Over the past decade, substantial efforts have been made to develop and understand evolution-based treatment strategies to combat drug resistance [5–21]. A key question in evolutionary biology is how likely and how quickly a resistant mutant will take over. While most research has focused on non-spatial or spatially homogeneous environments, an increasing number of recent theoretical and experimental studies have shown that spatial heterogeneity in drug concentration can significantly impact the evolutionary dynamics leading to resistance [22–53]. Intensive research has focused on the relationship between antibiotic resistance evolution speed and spatial drug heterogeneity. For example, it is well known that spatial drug heterogeneity can accelerate resistance evolution under specific spatial drug gradients [22–53]. Recent studies have also found that drug heterogeneity can slow down evolution [39–46]. Some authors argue that whether or not evolution is accelerated depends on spatial drug arrangements [36, 47, 48] and migration [49–53].

While evolution speed or emergence time is crucial for extending therapeutic windows and finding better resistance control strategies, another important quantitative feature, the phenotypic diversity of bacteria, should also be taken into account. Diversity quantifies a biological system’s ability of resilience and stability. Phenotypic diversity, such as a population with individual bacteria of varying levels of resistance, allows some fractions to survive environmental stress like antibiotic exposure. Investigating such phenotypic diversity in antibiotic resistance is essential because it underpins the ability of bacterial populations to adapt to and survive antibiotic treatments. This diversity makes treatment outcomes unpredictable and complicates efforts to eradicate bacterial infections [54, 55]. Moreover, migration between bacterial populations—whether across different physical locations or ecological niches—plays a key role in modulating phenotypic diversity. Migration introduces genetic and phenotypic variability, replenishes resistant individuals, and can distribute antibiotic-resistant traits more widely [24, 29, 56].

By tuning the level of migration, it is possible to influence how antibiotic resistance evolves within a population, either by promoting the spread of advantageous resistance phenotypes or by disrupting local adaptation through the mixing of less-resistant individuals. Therefore, understanding the role of migration is critical for designing strategies that can manage or mitigate the evolution of antibiotic resistance.

Recent research has shown that within-host diversity under antibiotic treatment is common [57], and within-patient diversity of pathogenic strain populations can accelerate antibiotic resistance evolution [58]. Investigating how spatial drug heterogeneity and migration modulate phenotypic diversity under antibiotic resistance may shed light on the existence of such diversity in human organs, subsequent mutations, or longer-term antibiotic resistance evolution. However, even in the simplest experimental heterogeneous environment with one drug-free sanctuary well and one drug well, systematic conclusions on how migration modulates bacterial diversity under drug resistance evolution are still lacking. The only evidence comes from a recent study that showed migration can delay but not prevent the spread of resistance at a fixed intermediate dispersal rate [40]. A well-designed 2-well migration-evolution experimental system is needed to fulfill the research needs in this field.

Although the diversity-migration relationship under antibiotic resistance evolution was previously unknown, evolutionary biologists have proposed a well-known relationship called the “Intermediate Disturbance Hypothesis” (IDH). The IDH suggests that disturbances, such as invader immigration and constant removal of local species, will induce maximum species diversity when the disturbance frequency is at an intermediate level [59]. However, despite its popularity, the IDH is empirical and lacks both experimental and theoretical support. Fieldwork is challenging for collecting data, and the commonly used competition-colonization dynamic framework, although it considers different spatial sites and spatial heterogeneities of selection pressure, only accounts for site occupation and lacks detailed microscopic dynamics of growth, competition, and migration [60]. A minimal model describing microscopic dynamics, considering growth and migration, is needed to quantitatively capture the diversity-migration relationship. A control parameter is also required to explain the different shapes of such relationships.

In this work, we designed a 2-well laboratory migration-evolution system with spatial drug asymmetry to mimic and generalize similar clinical bacterial behaviors, such as gut-lung translocation and antibiotic resistance evolution under different selection pressures [28]. Instead of focusing on resistance emergence speed, as has been extensively discussed in previous literature, we investigated how spatial drug asymmetry and migration modulate the diversity of different mutant phenotypes. Specifically, considering migration and its feedback as a form of “disturbance,” we found that, at the minimum inhibitory drug concentration, the diversity-migration relationship follows the “Intermediate Disturbance Hypothesis.” By tuning spatial drug asymmetry or, equivalently, the defined asymmetry parameter *k*, we experimentally revealed different diversity-migration relationships, which were validated by a minimal two-phenotype model describing microscopic growth-migration dynamics. Our findings were also tested with the bactericidal drug Ampicillin, yielding similar results when discrepancies induced by time-delayed responses were ignored. For two wells with two different types of drugs, four phenotypes were identified, including a cross-resistant type. The asymmetry parameter *k* still successfully captured the shapes of diversity-migration relationships for different drug concentration pairs. Our generalist-specialist framework explained the rich diversity-migration relationship details in this two-drug evolution through global fitness advantages. Our findings reveal how migration, as a disturbance, modulates phenotypic diversity in antibiotic resistance evolution. This understanding may help us better control pathogens that translocate between different organs.

## Results

Recent clinical findings from a critically ill patient have shown that the gut acts as a reservoir for subsequent lung colonization of extitPseudomonas aeruginosa, particularly under antibiotic treatment [28] (see Figure 1A). Understanding this translocation is key for clinical implications of pathogen resistance emergence at different body sites. To mimic and generalize such gut-lung translocation under various selection pressures, we designed a laboratory migration-evolution system using two wells in a 2 ml deep 96-well plate. Migration rates between the wells were controlled by mixing frequencies using the pipetting robot OT-2, with dilution at the start of each day, denoted as *β*. A 1:500 dilution was used to resume the exponential growth phase. The experiment was run for eight days to observe short-term bacterial resistance evolution (see more details in Supporting Information). The common gut bacterium extitE. faecalis was used. Since emigration removes local phenotypes and immigration introduces new invaders, we considered migration as a form of “disturbance” for each well. Our aim was to determine whether our diversity-migration relationship follows the well-known “Intermediate Disturbance Hypothesis” (IDH), where maximum diversity is expected at intermediate levels of disturbance. In our system, this translates to whether intermediate migration rates induce the highest phenotypic diversity.

**Figure 1.**
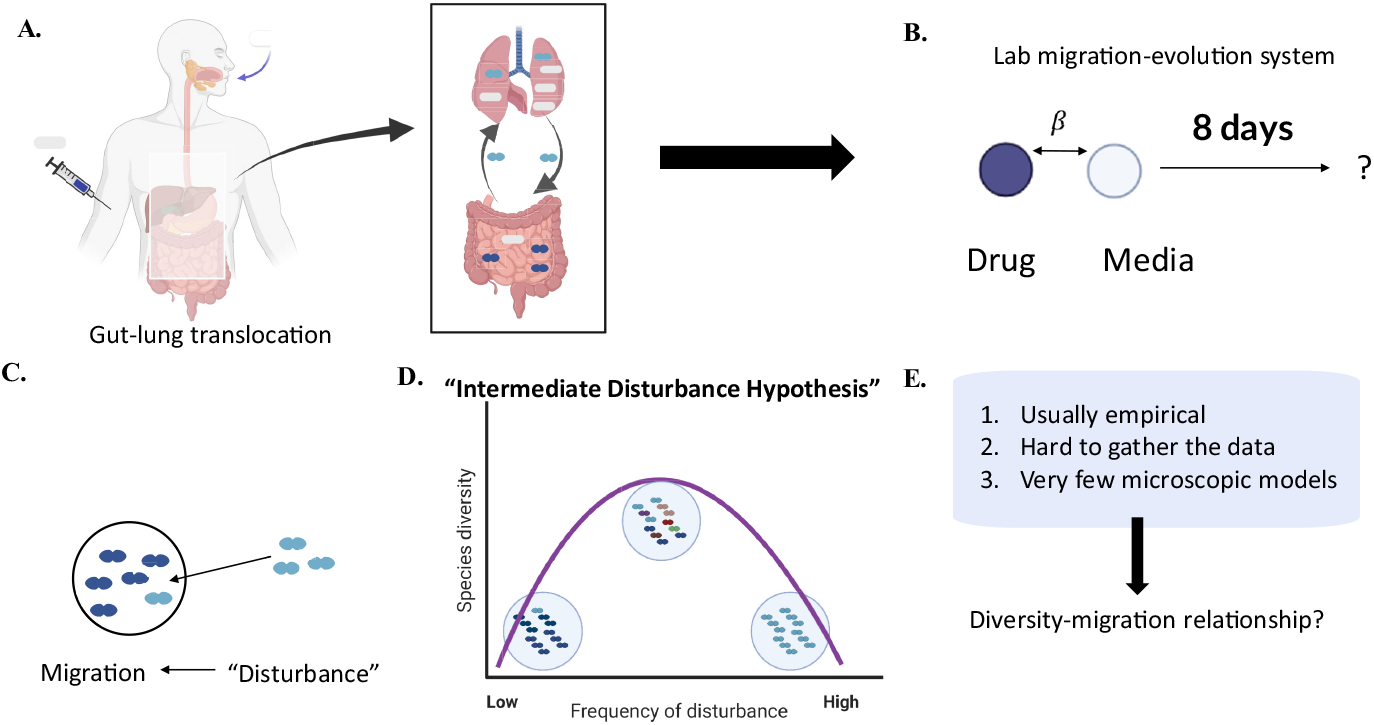
A schematic illustration of the lab migration-evolution system and the Intermediate Disturbance Hypothesis. **A, B**. An example of gut-lung translocation under different drug concentrations by PD/PK dynamics, and the simplified lab migration-evolution system. **C, D, F**. Treating migration as a disturbance, our phenotypic-diversity relationship may satisfy the “Intermediate Disturbance Hypothesis,” which is challenging to validate.

Understanding this relationship can facilitate the development of optimal clinical strategies for pathogen control. Specifically, intermediate migration levels may prevent highly resistant strains from dominating, while still maintaining microbial competition and diversity that can reduce complete colonization by resistant strains. This balance could be crucial in preventing outbreaks of highly resistant infections and in designing treatment protocols that use migration as a tool to manage microbial communities effectively. Additionally, IDH is challenging to validate in field studies, and mechanistic models are rare. Our experimental system provides a new approach to validating IDH and offers a model that captures explainable growth-migration microscopic dynamics.

To determine bacterial phenotypes, population-level samples were collected at the end of the experiments, and IC50 values (resistance levels) were determined by fitting drug dose-response curves. For the mono-antibiotic scenario, an IC50 threshold was used to classify populations as either sensitive or resistant. The proportions of each phenotype were obtained, and diversity was calculated using an appropriate metric. Although simple, this approach provides a minimal setting to explore the unknown diversity-migration relationship (see Figure 2).

**Figure 2.**
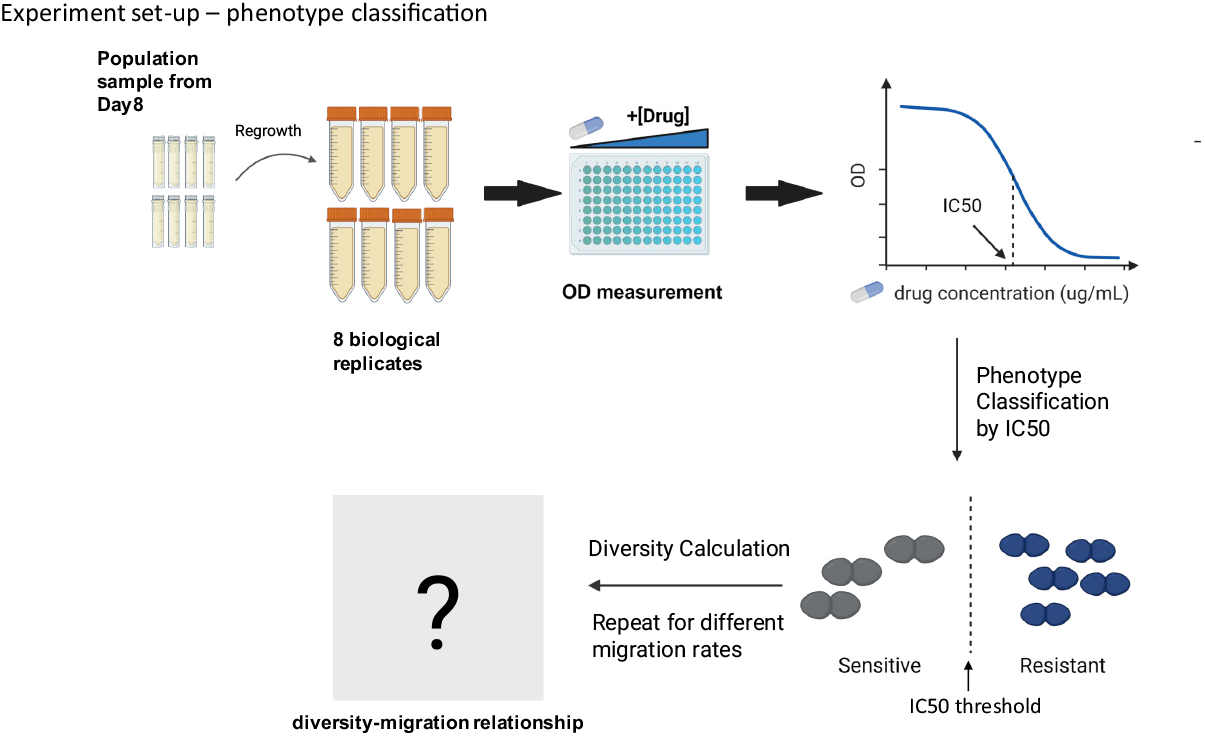
Experimental procedure for resistance level collection and pheno-type classification. Population samples from day 8 were collected and regrown for OD measurements. By classifying sensitivity and resistance based on fitted IC50 values, the diversity value was calculated for each migration level.

### Diversity-migration relationship satisfies “Intermediate Disturbance Hypothesis” under MIC

To create spatial drug asymmetry, we used the bacteriostatic drug Linezolid at the minimum inhibitory concentration (MIC) in the drug well. This drug concentration lies within the selection-mutation window, facilitating the emergence of drug-resistant phenotypes [61]. The other well contained only media. We chose Simpson diversity as our diversity metric, denoted as *H*, which also represents the “effective” number of phenotypes (see Figure 3B and Supporting Information). By modulating the migration rate *β*, the diversity-migration relationship showed a peak at the intermediate migration rate for both wells, satisfying the “Intermediate Disturbance Hypothesis.” The diversity in both wells at no migration (*β* = 0) and at relatively high migration rates (*β* = 5, 10) was 1 (see Figure 3C, left panel).

**Figure 3.**
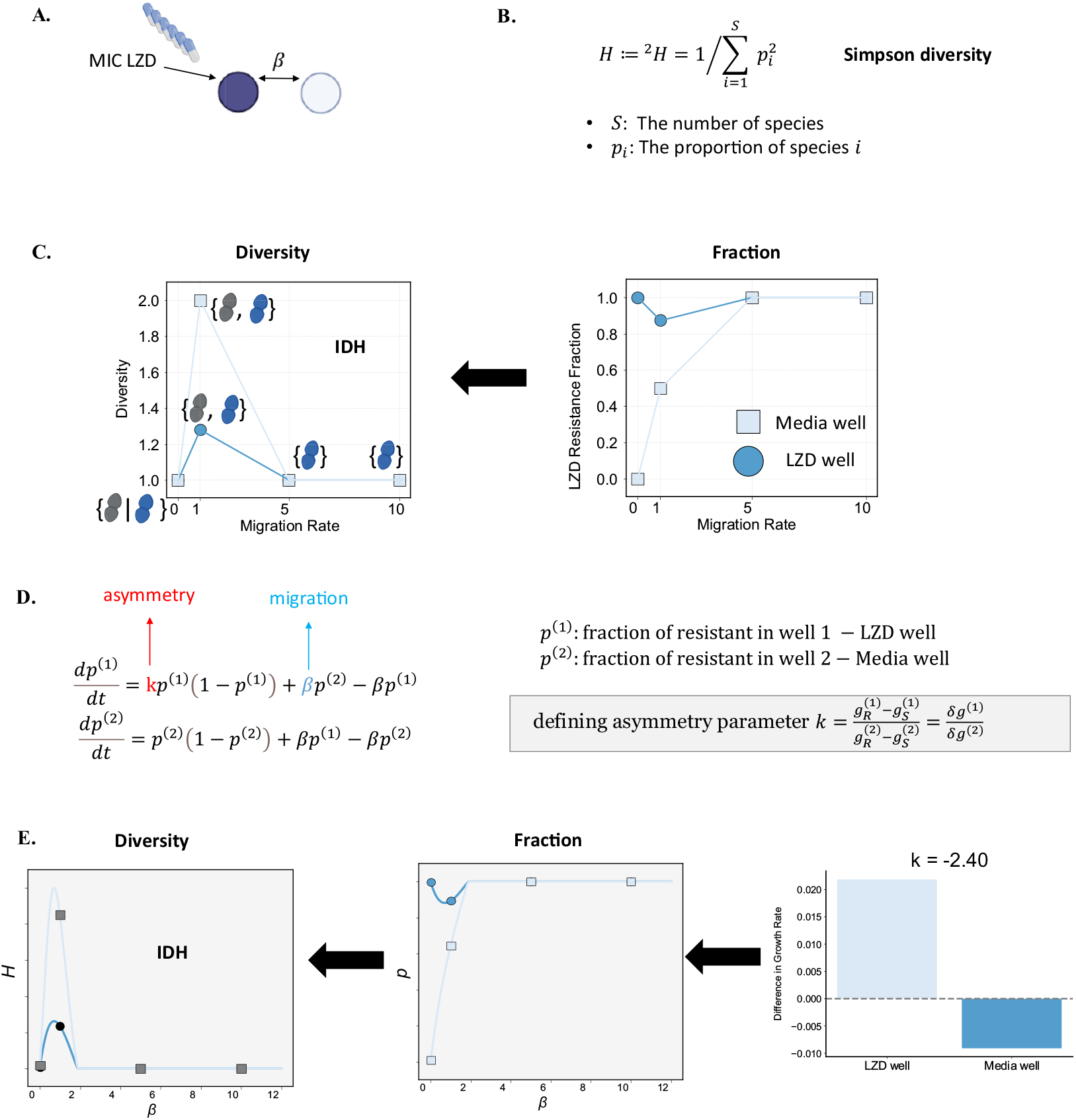
Diversity-migration relationship in both wells satisfies the “Inter-mediate Disturbance Hypothesis” under a specific spatial drug asymmetry. **A**. Spatial drug asymmetry with a minimum inhibitory concentration of Linezolid in the drug well. **B**. Simpson diversity is chosen as the diversity metric, representing the effective phenotype number. **C, D, E**. The experimental diversity-migration relationship satisfies IDH and can be explained by the phenotype fraction results. A minimal two-phenotype model introduces the single asymmetry parameter *k*, which controls the shape of the diversity-migration relationship. By using the experimentally determined value of this parameter in the model, we observed similar results in theory.

How do we explain this IDH pattern and the specific diversity values? This can be intuitively explained by examining the phenotype fractions. From the right panel of Figure 3C, we see that when there is no migration, only Linezolid-resistant phenotypes are present in the Linezolid well, and only sensitive phenotypes are present in the media well. This is expected, as local dominant phenotypes remain dominant without disturbance. Since diversity effectively represents the number of phenotypes, a diversity of 1 here indicates that there is only one phenotype in each well. When migration is introduced, the fraction of Linezolid-resistant bacteria in the Linezolid well decreases due to disturbance from the incoming invader and removal of resistant phenotypes from the local environment. Similarly, the fraction of Linezolid-sensitive bacteria decreases in the media well, but with a higher fraction change. This 50-50 fraction results in an effective phenotype number of 2, as reflected in the diversity panel, which is higher than the diversity in the Linezolid well. At migration rates of 5 and 10, Linezolid-resistant bacteria completely dominate both the Linezolid and media wells, showing a global fitness advantage, and the diversity returns to 1. The cartoon of extitE. faecalis in the left panel of Figure 3C also illustrates the change in phenotypes in each well.

To better understand this phenomenon, we present a minimal two-phenotype model describing the growth-migration dynamics in the two wells (see Figure 3D; for more details on model derivation, see Supporting Information):

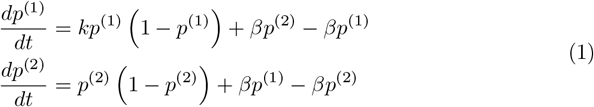

where *p*^(1)^ is the fraction of resistant bacteria in well 1 (the Linezolid well), and *p*^(2)^ is the fraction of resistant bacteria in well 2 (the media well). *k* is the asymmetry parameter defined as

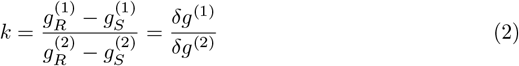

It represents the ratio of growth rate differences between the two wells (as a special case, *k >* 0 here also implies that one of the two phenotypes has a global advantage; see more details in Supporting Information). *β* is the migration rate, as defined earlier.

For this model, there are only two parameters: *β* and *k*. Since *β* already contributes to the diversity-migration relationship, *k* is the only parameter controlling the shape of this relationship. Thus, knowing *k* allows us to predict the shape of the diversity-migration relationship for this minimal two-phenotype model. By experimentally measuring the growth rate differences, we calculated *k* = −2.4 and used it in the model. The model-predicted fractions and diversities matched the experimental results well (see Figure 3C, E). We conclude that our diversity-migration relationship satisfies IDH under specific spatial drug asymmetry, controlled by a single asymmetry parameter *k*.

### Shape of diversity-migration relationships is determined by asymmetry parameter *k*

Since the asymmetry parameter *k* controls the shape of the diversity-migration relationship and is defined by growth rate differences, we adjusted the Linezolid concentrations in the drug well to modify these growth rate differences and observe whether relationships other than IDH might emerge. Six different concentrations, 1/6, 2/6, 3/6, 4/6, 5/6, 6/6 of the wild-type MIC were chosen, and for simplicity, these are denoted as drug concentrations 1, 2, 3, 4, 5, and 6. Figure 4B shows the corresponding drug dose-response curves of the sensitive and resistant phenotypes, obtained by measuring the growth rate of each population sample at the end of the evolution experiment and fitting the data. Here, we assumed that the resistant phenotypes emerging at different concentrations were similar (see more details in Supporting Information). Figure 4C,E shows the phenotype fractions and diversity results as migration rates change, based on experimental data. For higher drug concentrations (3, 4, 5, 6), we observed trends similar to those seen previously, with diversity-migration relationships satisfying IDH. However, for lower drug concentrations (1 and 2), there were no IDHs, and all diversity values were 1. This can be explained by examining the dose-response curves in Figure 4B. At low drug concentrations, the resistant phenotype had a lower growth rate compared to the sensitive phenotype due to fitness costs, making it unlikely for the resistant phenotype to be selected at drug concentrations 1 and 2. Thus, only the sensitive phenotype persisted.

**Figure 4.**
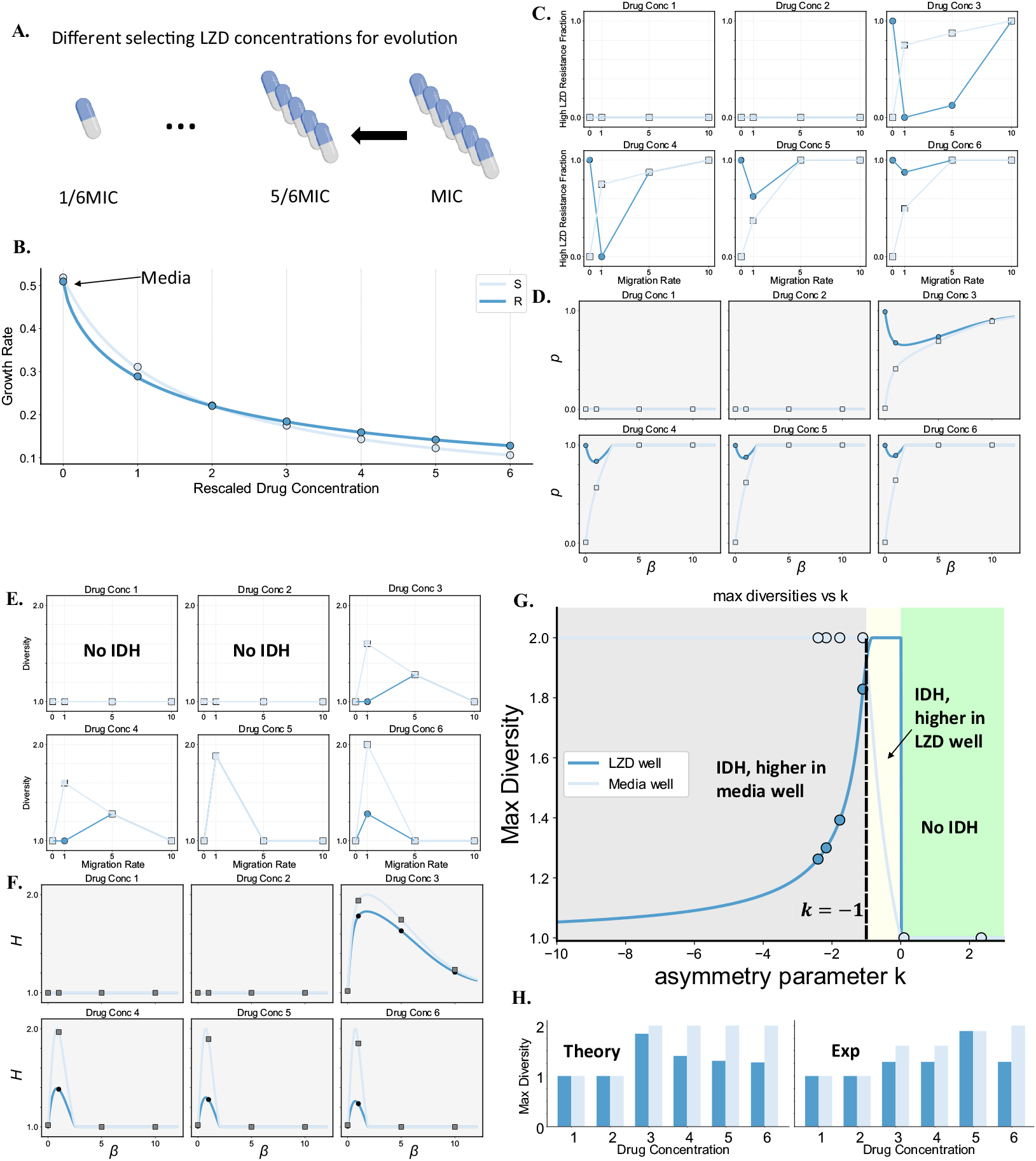
Different diversity-migration relationships emerge from different selecting Linezolid concentrations, explained by asymmetry parameter *k*. **A**. Six different Linezolid concentrations were selected. **B**. Corresponding dose-response curves for the sensitive and Linezolid-resistant phenotypes. **C, D**. Phenotype fractions as migration rate changes for six different Linezolid concentrations. The top panel shows experimental data, and the bottom panel shows theoretical simulations. **E, F**. Diversity-migration relationships for six different Linezolid concentrations. The top panel shows experimental data, and the bottom panel shows theoretical simulations. **G, H**. Phase diagram showing three different shapes of diversity-migration relationships, captured by maximum diversity and the asymmetry parameter *k*. Data points are positioned according to experimentally determined asymmetry parameters. Bar plots show the maximum diversity in both wells from theory and experimental data.

As mentioned earlier, the shape of the diversity-migration relationship is controlled by the asymmetry parameter *k* through growth differences. To quantify the relationship between shape and asymmetry, we chose the maximum diversity as a key quantitative feature. This approach allowed us to identify three distinct phases for the diversity-migration relationship as we tuned *k*, shown in Figure 4G. When *k <* −1, the diversity-migration relationship satisfies IDH, but with a higher diversity peak in the media well. In this case, the fraction in the media well is more sensitive to changes in migration. Drug concentrations 3, 4, 5, and 6 fall within this region, as indicated by the data points, and *k* values were determined experimentally. At the critical point *k* = −1, the maximum diversity values are equal in both wells, as the growth rates in the two wells are perfectly balanced. When −1 *< k <* 0, the diversity-migration relationship still satisfies IDH, but now the diversity peak is higher in the Linezolid well. When *k >* 0, there is no IDH, as one phenotype always dominates both wells, resulting in a maximum diversity of only 1, as seen in the two low drug concentration cases. Figure 4H shows the maximum diversity in both wells from both theoretical predictions and experimental data. Despite some discrepancies, the relative relationships of the maximum diversities were consistent between theory and experiment. These findings illustrate how different selecting Linezolid concentrations can lead to varying diversity-migration relationships, explained by the asymmetry parameter *k* in the phase diagram of maximum diversity.

### Bactericidal drug induces derivations from IDH while generally satisfies model predictions

To further validate and generalize our findings, we applied a bactericidal drug, Ampicillin (AMP), also at six different selecting drug concentrations. While bacteriostatic drugs like Linezolid inhibit bacterial growth, Ampicillin causes bacterial lysis, leading to population fluctuations and stochastic clearance of bacteria [62]. These features may alter diversity-migration relationships, as our minimal two-phenotype model only considers deterministic scenarios. Moreover, beta-lactam antibiotics like Ampicillin can lead to persistence or tolerance [63–66], inducing complex and potentially unknown dynamics such as time-delay responses or lag phases [67, 68], as well as the Eagle effect, characterized by a reduced lysis rate at high drug concentrations [69, 70]. Despite these complexities, we demonstrate that our deterministic framework still captures the qualitative shape of the diversity-migration relationship for Ampicillin.

Similar to the Linezolid scenario, Figure 5 shows how different diversity-migration relationships emerge for different selecting concentrations of Ampicillin. The dose-response curves of AMP-sensitive and AMP-resistant phenotypes also indicate the fitness cost for resistance at low drug concentrations (1 and 2), and a growth advantage at higher drug concentrations (3, 4, 5, 6) (see Figure 5B). Notably, the fraction-migration results for drug concentration 2 show a reversed trend compared to higher concentrations, in both experiments and theory (see Figure 5C, D, middle panels). This trend is indicated by a higher peak in the drug well, as confirmed by the phase diagram (Figure 5G, yellow region) and the theoretical diversity-migration relationship (Figure 5F, drug concentration 2). Unfortunately, this trend was not observed in the experimental diversity-migration relationship, likely due to the migration rates chosen.

**Figure 5.**
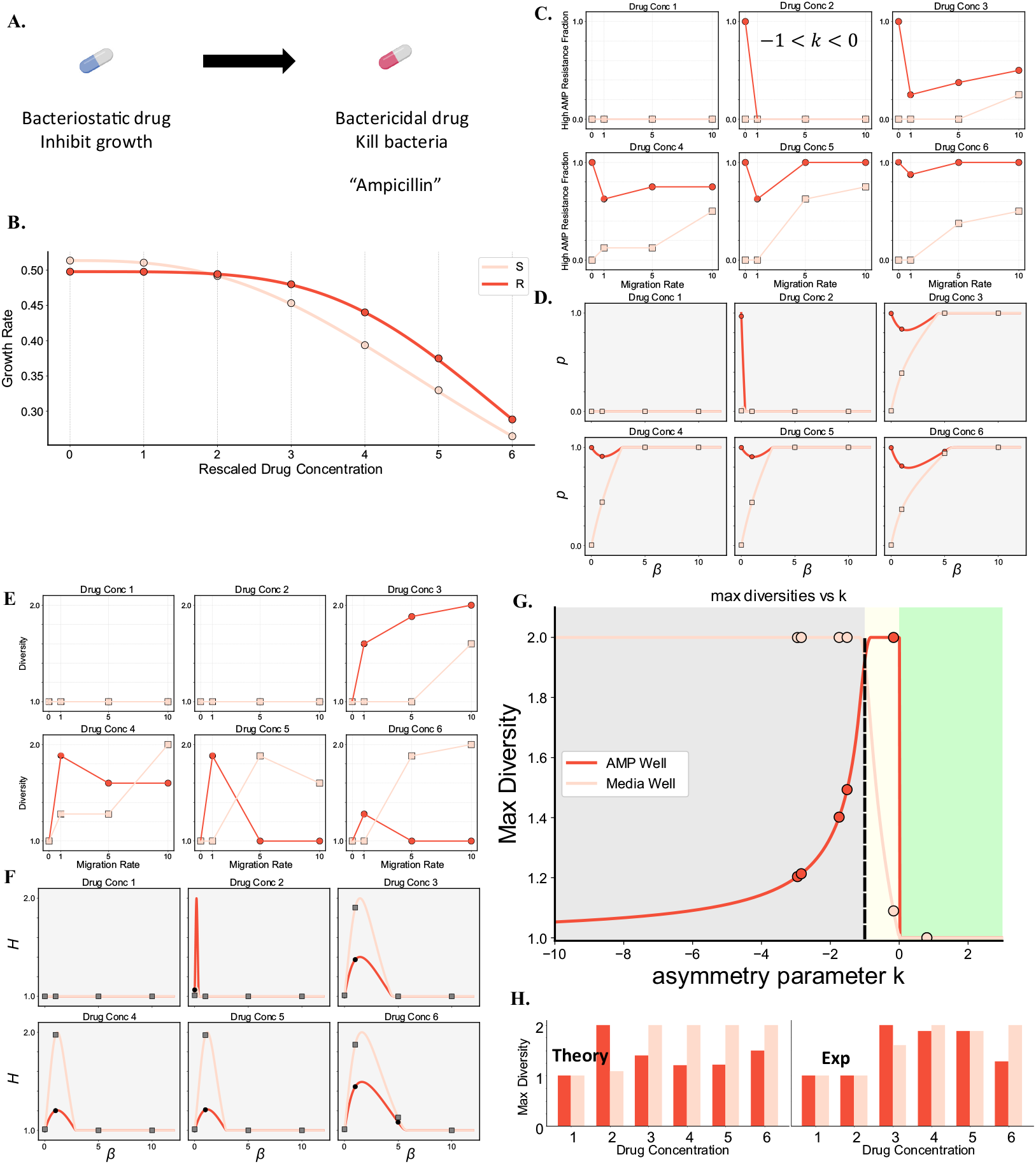
Different diversity-migration relationships emerge from different selecting concentrations of bactericidal drug Ampicillin, explained by asymmetry parameter *k*. **A**. Six different selecting drug concentrations were chosen. **B**. Corresponding dose-response curves for sensitive and AMP-resistant phenotypes. **C, D**. Phenotype fractions as migration rate changes for six different Ampicillin concentrations. The top panel shows experimental data, and the bottom panel shows theoretical simulations. **E, F**. Diversity-migration relationships for six different Ampicillin concentrations. The top panel shows experimental data, and the bottom panel shows theoretical simulations. **G, H**. Phase diagram showing three different shapes of diversity-migration relationships, captured by maximum diversity and asymmetry parameter *k*. Data points are positioned according to experimental asymmetry parameters. Bar plots show the maximum diversity in both wells, from theory and experimental data.

Although the fraction-migration results from simulations and experiments qualitatively match well (see Figure 5C, D), there is a time delay, or effectively lower migration rate, for the curves in the media well at high concentrations (3, 4, 5, 6). This is also evident in the experimental diversity-migration relationship (see Figure 5E, F). Furthermore, the maximum diversities for both wells do not emerge at the same migration rate as observed in the Linezolid case, with the relationship in the media well exhibiting a “slower” response. The simulation curves are still similar to those observed in the Linezolid case. All these observations indicate fundamental differences between bacteriostatic and bactericidal drugs, and evidence of bacterial tolerance was found through growth-death measurements (see more details in Supporting Information).

While more complex models could be used to explain these features, our minimal model still qualitatively captures the observed dynamics, even under more complex conditions. The comparison of maximum diversity between our model and experimental data also supports this qualitative match, with the only large discrepancy occurring at drug concentration 3 (see Figure 5H). Further detailed experiments could provide deeper insights into the complex dynamics observed. Nevertheless, our findings highlight the robustness of the minimal model and underscore the utility of the single shape-controlling asymmetry parameter *k* in capturing the diversity-migration relationship.

### Bacteriostatic-bactericidal spatial drug asymmetry drives similar diversity-migration relationships, and the shape is still captured by asymmetry parameter *k*

To demonstrate how spatial drug asymmetry and migration modulate diversity, we started with a relatively simpler 2-well system with one drug well and another as a sanctuary well. Now, we introduce a more complex scenario: a two-drug case where the bacteriostatic drug Linezolid is added to one well while the bactericidal drug Ampicillin is added to the other well. These are two common drugs used in first-line combination therapies for patients [71]. IC50 measurements for both drugs are necessary for phenotype classification. Besides the sensitive (S), Ampicillin-resistant (RA), and Linezolid-resistant (RL) phenotypes, a new cross-resistant (RR) phenotype emerged, which was not observed in the single-drug scenario. To confirm that this cross-resistance was not present in the single-drug cases, IC50s for resistant from both cases were also measured, and no RR phenotype was detected (see more details in Supporting Information). To further confirm that the cross-resistant type is not a mixed population of RL and RA, we selected 32 single isolates from sampled population replicates, and IC50 results showed that most of them were pure phenotypes (see more details in Supporting Information).

We aimed to determine if the diversity-migration relationships remain consistent under this scenario and whether some of them could still be captured by our asymmetry parameter *k*. Here, we extend the definition of *k* by considering the growth rate differences between the local dominant species *i*^*^ in well 1 (Linezolid well) and the local dominant species *j*^*^ in well 2 (Ampicillin well). Thus,

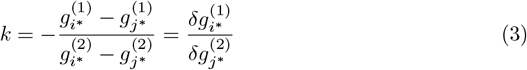

represents the ratio of growth rate differences between wells 1 and 2 (see Figure 6A; for more details, see Supporting Information).

**Figure 6.**
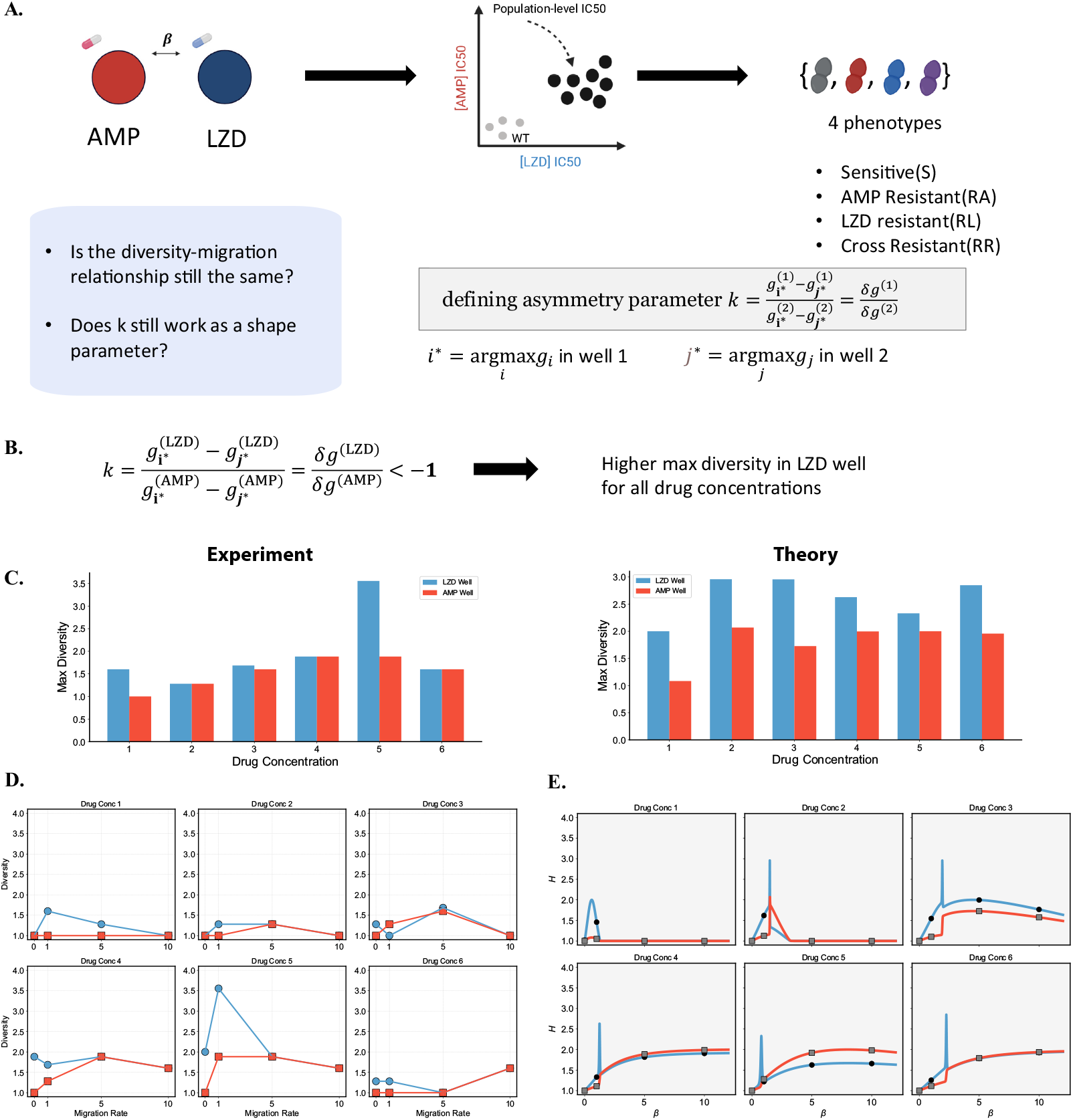
Spatial drug asymmetry by Ampicillin and Linezolid, and the diversity-migration relationships under six different selected drug pairs. **A**. Four different phenotypes—sensitive (S), Ampicillin-resistant (RA), Linezolid-resistant (RL), and cross-resistant (RR)—emerge from the migration-evolution experiments, according to the IC50 classification. The asymmetry parameter *k* is extended to incorporate more species. **B, C**. Experimental data calculated asymmetry parameter *k <* −1, indicating that higher maximum diversity occurs in the Linezolid well for all six different drug concentration pairs. Maximum diversity comparison in Linezolid and Ampicillin wells, from theory and experiment, confirms this. **D, E**. Diversity-migration relationships exhibit similar general shapes in experiment (Panel D) and theory (Panel E). Richer details and singular behaviors emerge in the theoretical predictions.

For all six selected drug concentration pairs, the asymmetry parameter *k* was found to be less than −1. This indicates that higher maximum diversity occurs in the Linezolid well for all drug concentrations, and the phenotypic fractions in the Linezolid well are more sensitive to changes in migration. This may still be explained by the possible tolerance in the Ampicillin well, which delays the response. As shown in Figure 6CDE, the diversity-migration relationship indeed exhibits higher peaks in the Linezolid well in both experimental and theoretical results. Although the general shapes appear similar, the theoretical predictions show more singular behaviors. This discrepancy arises due to the multi-phenotype nature of the system, and the fact that a single asymmetry parameter *k* can only capture the general shape, as it only reflects four growth rates. To account for richer details, an additional parameter would need to be introduced.

Despite the increased complexity, under different selected drug concentrations, the general shapes of the diversity-migration relationships in these complex systems are still captured by the single parameter *k*.

### Singular behaviors are explained by a minimal generalist-specialist framework and global fitness advantage

To explain the singular behavior and rich details in the simulation of the two-drug cases, we define a new parameter named the global fitness advantage:

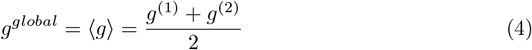

This parameter represents the spatial average of the growth rates in the two wells and is approximately the largest eigenvalue *λ*_1_ of our growth-migration system when the migration rate *β* is large (see derivations in Supporting Information). The phenotype with the largest *g*^*global*^ will dominate both wells at high migration rates. For example, in the two-phenotype model, when *k <* −1, this is equivalent to 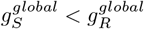, meaning the resistant phenotype will eventually dominate both wells, as shown in Figure 3.

We provide a minimal example involving three phenotypes—one generalist and two specialists—to explain the emergence of singular behaviors. The generalist has intermediate growth rates in both wells, while each specialist dominates one well with the highest growth rate there but has the lowest growth rate in the other well. Figure 7A shows that, by gradually increasing the global fitness advantage of the generalist 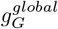, singular behaviors such as multiple peaks and crossings occur around the critical point, where

**Figure 7.**
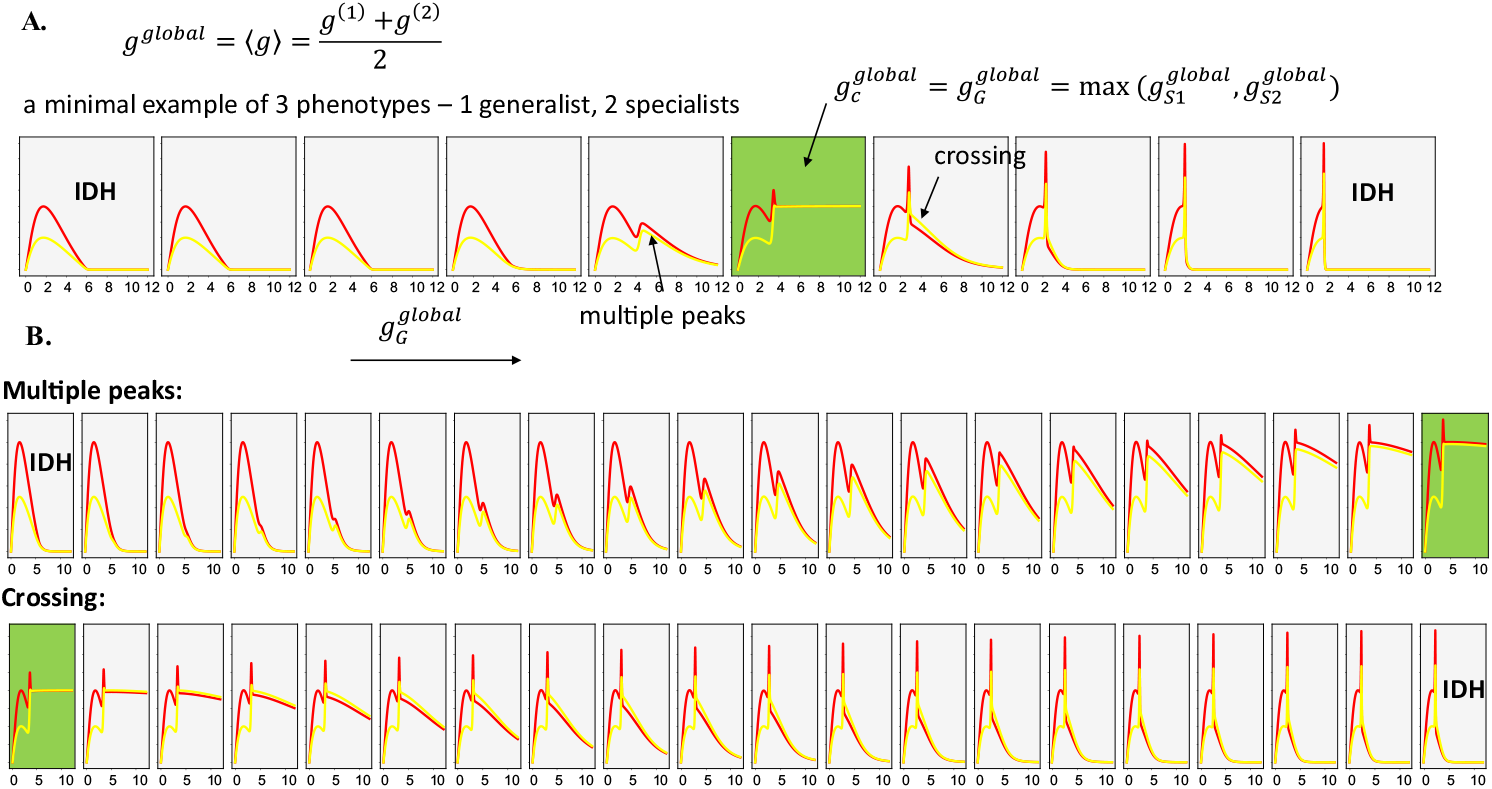
Global fitness advantages show how singular behaviors like multiple peaks emerge. **A**. A zoomed-in figure showing how IDH deforms into multiple peaks near the critical point 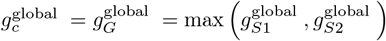. **B**. A zoomed-in figure showing how crossings return to the IDH near the critical point 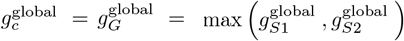.

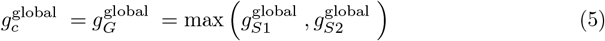

Beyond this critical point, the generalist becomes the globally dominant phenotype. If we zoom in, on the left side of the critical point, there is a narrow range of 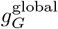 where the IDH gradually deforms into multiple peaks; on the right side of the critical point, there is also a narrow range of 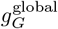 where the crossing gradually disappears and returns to a new IDH with sharp peaks. This provides an initial intuition as to why rich details and singular behaviors emerge. Since the parameter range of 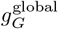 is very narrow and requires numerous migration rates to show a complete picture of interesting diversity-migration relationships, such as multiple peaks—a generalized version of IDH supported by both experimental and theoretical research [59, 72–77]—it is challenging to investigate experimentally using current lab techniques. However, it would be an intriguing area of study, as it could provide deeper insights into diversity-disturbance relationships in antibiotic resistance evolution and broader evolutionary dynamics.

## Discussion

In this work, we developed a two-well laboratory migration-evolution system that incorporates spatial drug asymmetry to systematically investigate how migration and drug gradients affect phenotypic diversity in bacterial populations. This setup serves to mimic and generalize clinical behaviors of bacteria, such as gut-lung translocation and the evolution of antibiotic resistance under different selection pressures [28]. Unlike most previous studies that focused heavily on the speed of resistance emergence, our primary focus was on how spatial drug asymmetry and migration influence the diversity of different mutant phenotypes—a factor critical for understanding pathogen control and devising optimal treatment strategies. By treating migration as a form of “disturbance” and working under minimum inhibitory concentrations, we found that our diversity-migration relationship aligns with the “Intermediate Disturbance Hypothesis” (IDH), which is a foundational but largely empirical concept in evolutionary biology. Through manipulating the spatial drug asymmetry or equivalently using our defined asymmetry parameter *k*, we experimentally revealed diverse diversity-migration relationships, which were validated using a minimal two-phenotype model. Testing our findings with Ampicillin, a bactericidal drug, showed similar results, albeit with discrepancies likely due to time-delayed responses. Further, in a two-drug scenario with Linezolid and Ampicillin, we identified four distinct phenotypes, including a cross-resistant type, and found that the asymmetry parameter *k* continued to effectively capture the diversity-migration relationships. Our generalist-specialist framework, based on global fitness advantage, helps explain the complex and emergent patterns of diversity-migration relationships observed in these more intricate settings. These findings contribute to a deeper understanding of how migration modulates phenotypic diversity during antibiotic resistance evolution, potentially offering new insights into pathogen control across different body sites.

Intensive research has focused on understanding the relationship between antibiotic resistance evolution speed and spatial drug heterogeneity. It is well-established that spatial drug heterogeneity can accelerate resistance evolution under specific spatial drug gradients [22–38]. Conversely, other studies have found that drug heterogeneity can also decelerate resistance evolution under different conditions [39–46]. Some researchers have even argued that whether evolution is accelerated or decelerated depends on the specific spatial arrangement of the drug and the extent of migration [36, 47–53]. While resistance speed is critical for extending the treatment window and finding effective control strategies, phenotypic diversity within bacterial populations also plays an essential role. Research shows that within-host diversity is common during antibiotic treatment [57], and within-patient diversity can accelerate resistance evolution [58]. Our study contributes to this body of work by focusing on short-term phenotypic diversity, offering new insights into how such diversity could exist within different organs of the human body and potentially influence the longer-term evolution of antibiotic resistance.

Spatial heterogeneity in selection pressures across different habitats has long been suggested as a mechanism for maintaining genetic variation in nature. Theories have proposed that migration, coupled with spatial heterogeneity, helps shape species diversity. For example, intermediate migration rates have been shown to enable gradual evolutionary diversification without requiring niches or tradeoffs [78]. In our study, we have demonstrated how even a simple two-well scenario with two different selection pressures can reveal the complexity of biodiversity dynamics. One-way migration, for example, has been shown to induce dynamical phase transitions and affect the time required to reach steady states [79]. Our results using a two-well system with automation platforms offer a unique experimental approach to further validate these theoretical predictions. The introduction of a single asymmetry parameter *k* as a metric to describe spatial growth asymmetry and predict the shape of diversity-migration relationships is, to our knowledge, the first of its kind, providing a straightforward way to connect spatial asymmetry with phenotypic diversity.

Although diversity-migration relationships in the context of antibiotic resistance were previously uncharted, evolutionary biologists have proposed the “Intermediate Disturbance Hypothesis” to describe how intermediate levels of disturbance, such as migration or environmental changes, can maximize species diversity [59]. Despite being well-known, IDH lacks experimental and theoretical support due to the challenges in field data collection and the absence of detailed mechanistic models. Our findings bridge this gap by offering a new method to investigate IDH, not only by generating quantitative data but also by providing a model that incorporates the underlying microscopic growth and migration dynamics. Our work suggests that the concept of IDH can be rigorously studied in controlled lab environments and extended to systems with antibiotic resistance evolution.

The emergence of multiple diversity peaks, which we observed under certain conditions, has also been documented in previous theoretical and experimental studies [72–76]. More recently, researchers found that multiple peaks can emerge within a generalized competition-colonization framework [77]. Our generalist-specialist framework offers a simple and intuitive explanation for this phenomenon by linking multimodal diversity-disturbance relationships to generalist-specialist dynamics. This connection provides a broader understanding of how species with different strategies interact to shape overall diversity in a system, and our minimal model captures these dynamics without requiring additional complex parameters.

While our experimental setup has provided significant insights, it is essential to note some limitations. This study only examined sub-lethal drug concentrations, which may not fully capture the diversity dynamics under higher, bactericidal concentrations. For bactericidal drugs like Ampicillin, bacterial lysis could lead to population clearance before resistance can develop, thereby altering phenotypic diversity outcomes [62].

Additionally, our system focused on a maximum of four phenotypes, whereas introducing more phenotypes could test the robustness of our asymmetry parameter and extend model predictions. Expanding the number of spatial sites could also help model more realistic networks, such as lymph nodes in cancer metastasis [80]. Moreover, we assumed symmetric migration between wells. Recent studies show that asymmetric migration, as seen in clinical settings like gut flow or metastasis, can decrease stability but increase resilience in microbial populations [81]. Exploring asymmetric migration experimentally could yield new insights into how migration shapes phenotypic diversity.

From a modeling perspective, we chose to ignore ecological interactions, assuming non-interacting populations due to daily dilution cycles. In natural settings, such interactions could include protective effects from resistant strains [82–84] or competition near carrying capacity [85–88]. Despite these simplifications, our minimal growth-migration model showed good agreement with experimental data, making it a valuable baseline for further studies, whether to explore specific system features or as a null model to detect new emergent properties. For spatial models considering ecological interactions, coexistence in competitive metacommunities is also found to be stable under environmental heterogeneities [89].Furthermore, we did not consider features specific to certain drugs, such as persistence, tolerance, or time-delayed responses.

Adding these features could help investigate, for example, how tolerance impacts diversity-migration relationships. Finally, our findings indicate that complex behaviors, such as those near critical points, require further experimental validation. These near-critical behaviors may be common in human systems and have been observed in other weakly ordered populations, such as brain cancer cells [90].

Our results raise several fundamental questions at the intersection of clinical, evolutionary, and basic microbiology. For example, how can we leverage maximum diversity to manage pathogen-related diseases? How can we design dynamic spatial drug asymmetries to adaptively control resistance evolution while maintaining bacterial populations within a manageable range, similar to adaptive therapy approaches [48, 91, 92]? Can the asymmetry parameter *k* serve as a predictor of harmful diversity levels, enabling proactive clinical interventions? We hope that our work motivates ongoing research into the complex interplay between spatial drug asymmetry, migration, optimal treatment strategies, and the evolutionary dynamics of resistance.

## Materials and Methods

### Strains, growth conditions, and drugs

Experiments were performed with *E. faecalis* strain OG1RF and the resistance was derived from the same strain. Samples were stocked in 20% glycerol and stored at -80°C. Cultures for experiments were taken from single colonies grown on agar plates and then inoculated at 37°C overnight before dilution in fresh media and the beginning of the experiment. All experiments and dose-response measurements were conducted in Brain-Heart Infusion or BHI (Remel). Drug stock solutions were prepared from powdered stock stored at -20°C and used in form of single-use aliquots stored at -80°C.

**Table 1.** Table of antibiotics used in this study and their targets.

| Drug Name (Abbreviation) | Supplier | Class | Mechanism of Action |
| --- | --- | --- | --- |
| Ampicillin Sodium Salt % (AMP) | Fisher Scientific | β-lactam (penicillins) | Cell wall synthesis inhibitor |
| Linezolid, 98% (LZD) | Acros Organics | Oxazolidinone | 50S protein synthesis inhibitor |

### Automated laboratory evolution experiments with migration

Evolution experiments in each pair of wells (drug and media, or AMP and LZD) were performed in 8 biological replicates, with Opentrons OT-2 pipetting robot doing the liquid transfer as migration. Drug concentrations at 1/6, 2/6, 3/6, 4/6, 5/6, 6/6 of WT MIC for AMP and LZD of WT were used, totally 6 different pairs. Evolutions were performed using 1ml BHI medium in 96-well plates with a maximum volume of 2ml. Each day, populations between 2 paired wells were mixed 0,1,5,10 times back and forth with a single-time 200ul volume transfer. A 1/500 dilution was used to inoculate the next day’s evolution plate, and the process was repeated for a total of 8 days of selection. On the final day of evolution, all strains were stocked in 20% glycerol. Strains were then streaked from the population-level samples for population-level IC_50_ determination. In the case of AMP-LZD evolution, days 1, and 4 were also stocked for further testing, and single isolate level IC50s were also determined by plating selected strains on a pure BHI plate, and selecting 32 single colonies for each strain.

### Phenotypic resistance profiling - measuring drug resistance and sensitivity

Experiments to estimate IC_50_ were performed in 96 -well plates for both population-level and single-isolate level strains. by exposing mutants to a drug gradient consisting of 12 points-one per well—typically in a linear dilution series prepared in BHI medium with a total volume of 195 *µ*L(195*µ*L of BHI, 5*µ*L of cells) per well. After 16-20 hours of growth, the OD at 600nm (OD600) was measured using an Enspire Multimodal Plate Reader (Perkin Elmer) with an automated 20-plate stacker assembly. This process was repeated for all mutants with 8 replicates of WT as control for population-level strains and 64 replicates of WT for single-isolate level strains.

The OD (OD600) measurements for each drug concentration were normalized by the OD600 in the absence of drug. To quantify drug resistance, the resulting dose-response curve was fit to a Hill-like function *f* (*x*) = (1 + (*x/K*)^*h*^) ^−1^ using nonlinear least squares fitting, in which *K* is the *IC*_50_ and *h* is a Hill coefficient describing the steepness of the dose-response relationship. For drug-media evolution, A mutant strain was defined to be sensitive(S) if its IC_50_ smaller than the resisrance threshold, which is set to be at least larger than 3*σ*_*a*_ relative to the ancestral strain (3*σ*_*a*_ is defined as the uncertainty—standard error across replicates-of the IC_50_ measured in the ancestral strain). Similarly, an increase in IC_50_ by the threshold at least larger than 3*σ*_*a*_ relative to the ancestral strain corresponds to resistance(R). For AMP-LZD evolution, 2 resistance thresholds are used to divide mutants into 4 classes: S, RA, RL, RR, according to their IC50s. S is sensitive, RA is AMP-resistant, RL is LZD-resistant, and RR is cross(double) resistant to both drugs.

### Growth rate measurement

To reduce variance of experimental observation, per capita growth rate (g) from OD time series were estimated, by fitting the early exponential phase portion of the background subtracted curves before a time cutoff *T*, to an log-transformed time-derivative function (*d* ln(OD)*/dt* = *g*), using linear least-square fitting [93]. If the growth rate was estimated to be negative or the growth curve did not reach 0.001 over the course of the experiment, growth was set to 0. All growth rates were normalized by the growth rate of ancestral strains (WT) in the absence of drugs performed on the same day.

For growth rates at different selecting drug concentrations, all the strains were selected are measured. For growth dose response measurements of selected strains, 6 drug concentrations were selected the same as the selecting drug concentrations, and the resulting dose-response curve was fit to a Hill-like function *f* (*x*) = *g*_0_(1 + (*x/K*)^*h*^)^−1^ using nonlinear least squares fitting, in which *g*_0_ is drug-free growth rate, *K* is the *IC*_50_ and *h* is a Hill coefficient describing the steepness of the dose-response relationship.

### Tolerance measurement

Tolerance strains usually have no genetic change but a smaller growth rate and death rate [63, 65, 67, 94]. To find whether or not AMP-tolerant strains emerge from the sub-lethal AMP environment, drug-free growth rates and maximum death rate were estimated using the method from [93]. Overnight bacteria were diluted 100-fold before experiments. Strains were grown for 3-5hs and OD were measured by the Enspire Multimodal Plate Reader (Perkin Elmer) before adding AMP. Drug-free growth rates were estimated as above. After adding AMP, the rate of change in OD is a combination of continued cell growth and death due to the bactericidal feature of AMP:*dln*(*OD*)*/dt* = *g*− *d*. Based on previous observations, the instantaneous pre-antibiotic growth rate was assumed to remain constant [95, 96] after the addition of the antibiotic:*d* = *g* − (*dln*(*OD*)*/dt*). Death rate *d* is a function of time and maximum death rate *d*_*m*_ was chosen as a metric of the lysis dynamics. The tolerance level were determined by comparing *g* and *d* between mutants and WT(See Supporting Information). Strains collected at Day 8 were used to measure the population-level tolerance. From each of 10 population-level mutants, 32 single colonies were selected to measure the heterogeneous level of tolerance.

### Hill number as a diversity metric

Hill integrated species richness and species abundances(or proportions) into a class of diversity measures later called Hill numbers, or effective numbers of species [97, 98], defined for *q*≠ 1 as

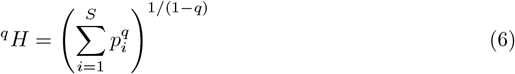

in which *S* is the number of species, and *p*_*i*_, *i* = 1, 2, …, *S* is the proportion of the *i* th species. The parameter *q* determines the sensitivity of the measure to the relative frequencies. At *q* = 0, the frequencies or proportions of species don’t contribute to the diversity and ^0^*H* is simply species richness(or absolute species number). For simplicity we chose *q* = 2, which yields a common used metric in evolutionary biology named as Simpson diversity:

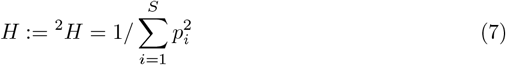

For convenience, in this paper *H* is used to denote Simpson diversity.

### Model simulation

To explain how migration modulates the evolution, we considered a simple mathematical model of exponentially growing subpopulations (sensitive and resistant cells) whose growth rates depend on the environment. All strains shared the same migration rate differing from each other in their growth rates, governed by the dose response curves at different selecting drug concentrations. All the simulations are initialized with the same initial condition, and integrated within [0, 24×8] h. It’s long enough to reach the steady state(Supporting Information). Migration rate is the known control parameter, and growth rates are estimated from OD curves, so there is no free parameter in our model. The simulation is performed using the *odeint* function from the Python package *SciPy*.

## Acknowledgments

This study was supported by NIH R35GM124875 (KBW).

## Notes

### Competing Interest Statement

The authors have declared no competing interest.

