## Supplementary Information for "Spatial drug asymmetry modulates phenotypic diversity-migration relationships under resistance evolution"

#### Supporting Information

This document contains the supporting information for the main manuscript titled "Spatial drug asymmetry modulates phenotypic diversity-migration relationships under resistance evolution". The sections below contain additional figures, methods, and data.

#### Intermediate Disturbance Hypothesis under antibiotic resistance evolution

In ecological and evolutionary dynamics, the Intermediate Disturbance Hypothesis (IDH) proposes that species diversity is maximized at intermediate levels of disturbance due to a balance between competitive exclusion and disturbance-driven population turnover[1]. In environments with low disturbance, strong competitors dominate and exclude weaker species, reducing overall diversity. Conversely, at high disturbance levels, few species can survive the constant disruptions. However, with intermediate disturbances, both competitive and resilient species coexist, fostering higher diversity.

Since IDH is mainly due to the tradeoff between competition and disturbance, usually typical the competition is due to different growth rates, and the disturbance is due to population removal, or possible invaders. In the case of proposed two-well lab system, if we are continuously mixing bacterial phenotypes between both wells, several key elements of the IDH are present: a. Competition by Growth Rates: The different bacterial species are in competition, and the competition rate is linked to their respective growth rates. This aligns with the aspect of competition that is central to the IDH. b. Disturbance via Mixing: The mixing action can be seen as a form of disturbance that prevents any single species from fully monopolizing either well. As long as the mixing is happening at an "intermediate level"—meaning not too frequent and not too rare—it should maintain a balance between competitive exclusion and allowing colonization opportunities for other species. This would align well with the conditions needed to study IDH.

### Methods

Detailed information about the methods used in this study can be found here.

#### Model-related methods and supplementary results

##### N-phenotype growth-migration dynamics over M patches, proportion dynamics, and diversity dynamics

A general N-phenotype growth-migration dynamics in M wells are described as

$$\begin{gathered}\frac{\partial u_i}{\partial t} = -\beta L \mathrm{u}_i + G_i(D) u_i \\ i = 1, 2, \cdots N\end{gathered} \tag{S1}$$

where

$$G_i = \begin{pmatrix} g_i^{(1)}(D) & & & \\ & g_i^{(2)}(D) & & \\ & & \ddots & \\ & & & g_i^{(M)}(D) \end{pmatrix}$$

is a diagonal growth matrix of species $i$, with each diagonal entry describing the growth rate controlled by drug concentration $D$ at different well positions. In the following demonstrations, we will omit writing $D$ for simplicity. $u_i$ is the population density of species $i$ over $M$ wells/patches. $\beta$ is the migration rate. $L$ is the Laplacian matrix describing the connectivity structure of wells. We can transform this dynamics of population density change to the dynamics describing change of phenotype proportions as

$$\dot{p}_i^{(m)} = p_i^{(m)} \left( g_i^{(m)} - \langle g^{(m)} \rangle \right) - \beta \sum_n L_{mn} \left( p_i^{(n)} - p_i^{(m)} \right) \tag{S2}$$

where $m$ denotes $m$th well, $\langle g^{(m)} \rangle = \sum_{i=1}^N p_i g_i^{(m)}$ and $\sum_{i=1}^N p_i^{(m)} = 1$. For 2 wells, this becomes

$$\begin{aligned}\dot{p}_i^{(1)} &= \delta g_i^{(1)} p_i^{(1)} - \beta \left( p_i^{(1)} - p_i^{(2)} \right) \\ \dot{p}_i^{(2)} &= \delta g_i^{(2)} p_i^{(2)} - \beta \left( p_i^{(2)} - p_i^{(1)} \right)\end{aligned} \tag{S3}$$

$i = 1, 2, \cdots N$. And $\delta g_i^{(m)} = g_i^{(m)} - \sum_{j=1}^N x_j^{(m)} g_j^{(m)}, m = 1, 2$

Recall we have Hill diversity defined as

$${}^qH = \left( \sum_{i=1}^S p_i^q \right)^{1/(1-q)}$$

At $q = 2$, we get the Simpson diversity

$${}^2H = 1 / \sum_{i=1}^S p_i^2 \tag{S4}$$

Do time derivative of ${}^2H$ for both wells and take advantage of the proportion dynamics by inserting $\dot{p}_i^{(1)}, \dot{p}_i^{(2)}$, after some re-arrangement we have the diverstiy dynamics

$$\begin{aligned} \dot{H}^{(1)} &= -2H^{(1)}\left[H^{(1)}\,\mathrm{Cov}_{p^{(1)}}\left(\delta g^{(1)}, p^{(1)}\right) - \beta + \beta H^{(1)} O\right] \\ \dot{H}^{(2)} &= -2H^{(2)}\left[H^{(2)}\,\mathrm{Cov}_{p^{(2)}}\left(\delta g^{(2)}, p^{(2)}\right) - \beta + \beta H^{(2)} O\right] \end{aligned} \tag{S5}$$

where $\mathrm{Cov}_{p^{(m)}}\left(\delta g^{(m)}, p^{(m)}\right) = \sum_{i=1}^{S} \delta g_i \left(p_i^{(m)}\right)^2, m = 1, 2$. $O = \sum p_i^{(1)} p_i^{(2)}$ is inverse of an overlapping diversity, or overlap coefficient, measuring the similarity between the species distributions of the two groups. We can thus simulate the diversity dynamics with the simple forms above(or we can still do the simulations by proportion dynamics since the time derivatives of Simpson diversity is not in closed form).

For $O$, we should have a simple relationship with diversities in 2 wells as $1/H^{(1)} + 1/H^{(2)} + 2O = 4$. Thus, we can also derive time derivative of the overlap diversity as

$$\begin{aligned} \dot{O} &= \sum_{i=1}^{S}\left(\delta g_i^{(1)} + \delta g_i^{(2)}\right) p_i^{(1)} p_i^{(2)} + \beta\left(\frac{1}{H^{(1)}} + \frac{1}{H^{(2)}} - 2O\right) \\ &= \langle p^{(1)} | \delta g^{(1)} + \delta g^{(2)} | p^{(2)} \rangle + \beta\left(\frac{1}{H^{(1)}} + \frac{1}{H^{(2)}} - 2O\right) \\ &= \langle p^{(1)} | \delta g^{(1)} + \delta g^{(2)} | p^{(2)} \rangle + 4\beta\,(1 - O) \end{aligned}$$

If rewrite the covariance between growth rate difference and the proportions as

$$\mathrm{Cov}_{p^{(m)}}\left(\delta g^{(m)}, p^{(m)}\right) = \frac{1}{H^{(m)}}\left(\langle g^{(m)}\rangle_{q^{(m)}} - \langle g^{(m)}\rangle_{p^{(m)}}\right)$$

where $q_i^{(m)} = \frac{\left(p_i^{(m)}\right)^2}{D^{(m)}}$ is a new probability distribution emphasizing more abundant species. Thus we can also rewrite the diversity dynamics as follows

$$\begin{aligned} \dot{H}^{(1)} &= -2H^{(1)}\left(\left\langle g^{(1)}\right\rangle_{q^{(1)}} - \left\langle g^{(1)}\right\rangle_{p^{(1)}} - \beta + \beta H^{(1)} O\right) \\ \dot{H}^{(2)} &= -2H^{(2)}\left(\left\langle g^{(2)}\right\rangle_{q^{(2)}} - \left\langle g^{(2)}\right\rangle_{p^{(2)}} - \beta + \beta H^{(2)} O\right) \end{aligned} \tag{S6}$$

Although it's not our main focus here, this introduction of distribution transformation may have interesting further interpretations.

##### 2-phenotype model in 2 well and asymmetry parameter $k$

In the experimental migration-evolution system, if we only apply drug to 1 well, the minimal resistance level classification gives us 2 phenotypes - sensitive and resistant. Here we give a detailed derivation of the simplest version of 2-phenotype model mentioned in main text, also introduce the asymmetry parameter $k$, for tuning the shape of the diversity-migration relationship. With the derived proportional dynamics in 2 well, if limiting the phenotype number to 2, it becomes 2 equations with reduced form

$$\begin{aligned}\frac{dp^{(1)}}{dt} &= \delta g^{(1)} p^{(1)} \left(1 - p^{(1)}\right) + \beta p^{(2)} - \beta p^{(1)} \\ \frac{dp^{(2)}}{dt} &= \delta g^{(2)} p^{(2)} \left(1 - p^{(2)}\right) + \beta p^{(1)} - \beta p^{(2)}\end{aligned}$$

where $p^{(1)}$ is the fraction of resistant in well 1, $p^{(2)}$ is the fraction of resistant in well 2, and $\delta g^{(1)} = g_R^{(1)} - g_S^{(1)}, \delta g^{(2)} = g_R^{(2)} - g_S^{(2)}$ are the growth rate differences between resistant and sensitive in 2 wells. Thus 2 equations here determine the whole dynamics instead of 2N equations for N phenotypes in 2 wells. Here we can define the asymmetry parameter $k = \frac{g_R^{(1)} - g_S^{(1)}}{g_R^{(2)} - g_S^{(2)}} = \frac{\delta g^{(1)}}{\delta g^{(2)}}$, to describe the spatial asymmetry level by growth rates in 2 wells. Define $\delta_g = \delta g^{(1)}$, and we get the following equations

$$\begin{aligned}\frac{dp^{(1)}}{dt} &= k \cdot \delta_g p^{(1)} \left(1 - p^{(1)}\right) + \beta p^{(2)} - \beta p^{(1)} \\ \frac{dp^{(2)}}{dt} &= \delta_g p^{(2)} \left(1 - p^{(2)}\right) + \beta p^{(1)} - \beta p^{(2)}\end{aligned}$$

Do the time rescaling and let $\delta_g = 1$ without loss of generality, and we have

$$\begin{aligned}\frac{dp^{(1)}}{dt} &= k p^{(1)} \left(1 - p^{(1)}\right) + \beta p^{(2)} - \beta p^{(1)} \\ \frac{dp^{(2)}}{dt} &= p^{(2)} \left(1 - p^{(2)}\right) + \beta p^{(1)} - \beta p^{(2)}\end{aligned} \tag{S7}$$

As mentioned in the main text, this equation system only has 2 effective parameters, with migration rate $\beta$ contributed to the diversity-migration relationship. Asymmetry parameter $k$ thus becomes the only controlling parameter modulating the shape of this relationship. Thus we have a minimal explainable model to interpret the experimental results. Different from the empirical models like colonization-competition dynamics, this is also a minimal microscopic dynamic system considering growth and migration, from what we know, to induce the behavior satisfying "Intermediate Disturbance Hypothesis".

We can similarly derive the diversity dynamics for 2 phenotypes

$$\begin{aligned}\dot{H}^{(1)} &= 2\left(1 - 2p^{(1)}\right)\left[k p^{(1)} \left(1 - p^{(1)}\right) + \beta \left(p^{(2)} - p^{(1)}\right)\right] \left(H^{(1)}\right)^2 \\ \dot{H}^{(2)} &= 2\left(1 - 2p^{(2)}\right)\left[p^{(2)} \left(1 - p^{(2)}\right) + \beta \left(p^{(1)} - p^{(2)}\right)\right] \left(H^{(2)}\right)^2\end{aligned} \tag{S8}$$

**A Padé analytical approximation for diversity-mgiration relationship $H(\beta)$**

Recall the 2-phenotype model

$$\begin{aligned}\frac{dp^{(1)}}{dt} &= -|k|p^{(1)}\left(1-p^{(1)}\right)+\beta p^{(2)}-\beta p^{(1)} \\ \frac{dp^{(2)}}{dt} &= p^{(2)}\left(1-p^{(2)}\right)+\beta p^{(1)}-\beta p^{(2)}\end{aligned} \tag{S9}$$

Since $k > 0$ only gives us diversity 1 as shown in main text. Here we only consider $k < 0$ and write $k = -|k|$ for simplicity. We also set $\delta_g = 1$.

At equilibrium $\left(\frac{dp^{(1)}}{dt} = \frac{dp^{(2)}}{dt} = 0\right)$, the equations become:

$$\begin{aligned}0 &= -|k|p^{(1)}\left(1-p^{(1)}\right)+\beta\left(p^{(2)}-p^{(1)}\right) \\ 0 &= p^{(2)}\left(1-p^{(2)}\right)+\beta\left(p^{(1)}-p^{(2)}\right)\end{aligned}$$

Alternatively, consider small deviations of resistant fractions: $p^{(1)} = 1-\delta p^{(1)}$ in well 1, and $p^{(2)} = \delta p^{(2)}$ in well 2. Plug into the equilibrium equations:

$$\begin{aligned}0 &= -|k|\left(1-\delta p^{(1)}\right)\left(\delta p^{(1)}\right)+\beta\left(\delta p^{(2)}-1+\delta p^{(1)}\right) \\ 0 &= \delta p^{(2)}(1-\delta p^{(2)})+\beta\left(1-\delta p^{(1)}-\delta p^{(2)}\right)\end{aligned}$$

Retain terms up to first order in $\delta p$ and neglect higher-order terms like $(\delta p)^2$ :

$$\begin{aligned}0 &\approx -|k|\delta p^{(1)}+\beta\left(1-\delta p^{(2)}\right)-\beta\delta p^{(1)} \\ 0 &\approx \delta p^{(2)}+\beta\delta p^{(1)}-\beta+\beta\delta p^{(2)}\end{aligned}$$

Doing the re-arrangement, we get the solutions

$$\begin{aligned}\delta p^{(1)} &\approx \frac{\beta}{|k|+\beta(|k|+1)} \\ \delta p^{(2)} &\approx \frac{|k|\beta}{|k|+\beta(|k|+1)}\end{aligned} \tag{S10}$$

From this result we can see that, $\delta p^{(2)} \approx |k|\delta p^{(1)}$. This explains when turning on the migration, the sensitivity of resistant fractions(or the fraction change) in media well is higher than that in drug well, as seen in main text. Thus the expressions for diversity-migration relationship $H^{(1)}(\beta)$ and $H^{(2)}(\beta)$ at equilibrium are:

$$\begin{aligned}H^{(1)}(\beta) &= \frac{1}{\left(1-\delta p^{(1)}\right)^2+\left(\delta p^{(1)}\right)^2} = \frac{[k+\beta(k+1)]^2}{\beta^2+[k(1+\beta)]^2} \\ H^{(2)}(\beta) &= \frac{1}{\left(1-\delta p^{(2)}\right)^2+\left(\delta p^{(2)}\right)^2} = \frac{[k+\beta(k+1)]^2}{(k+\beta)^2+(k\beta)^2}\end{aligned} \tag{S11}$$

However, this is only true at small $\beta$ - it will go to a constant as migration rate increases. Besides to explain how $k$ tunes the general changes, We also need approximated diversities to satisfy different phases with different $k$s, and goes back to 1 as $\beta \to \infty$.

$H(\beta)$ **expressions as Padé approximants** By observation, our approximated diversities have the form of [2/2] Padé approximants[2, 3]. We aim to reconstruct a [1/n] Padé Approximant so dversity goes back to 1 as $\beta \to \infty$. We can use a [1/2] Padé Approximant in the form $H(\beta) = 1 + \frac{A\beta}{1+B\beta+C\beta^2}$ as a minimal example, so that this new approximant satisfies following conditions: (a) At $\beta = 0, H(\beta) = 1$; (b) As $\beta \to \infty, H(\beta) \to 1$; (c) $k = 1$, $H^{(1)}(\beta) = H^{(2)}(\beta)$, ensuring symmetry; (d) When $k \neq 1$, $H^{(1)}(\beta) \neq H^{(2)}(\beta)$, ensuring asymmetry. Let's match the coefficients of taylor series of the original diversity function at $\beta = 0$, with of [1/2] Padé Approximants to the third order $\beta^3$ ([m/n] Padé approximant requires matching coefficients with original function $f(x)$ to the highest possible order $f^{(m+n)}(0)$). Supposing the small-$\beta$ expansion of original $H(\beta)$ is $H(\beta) = 1 + h_1\beta + h_2\beta^2 + h_3\beta^3 + \ldots$, we have

$$\begin{aligned} H^{(1)}(\beta) &= 1 + \frac{2}{k}\beta + \left(-2 - \frac{2}{k} + \frac{2}{k^2}\right)\beta^2 + \left(-k^2 - \frac{8}{3}k - \frac{4}{3}\right)\beta^3 + \mathcal{O}\left(\beta^4\right) \\ H^{(2)}(\beta) &= 1 + 2\beta + \left(2 - \frac{2}{k} - \frac{2}{k^2}\right)\beta^2 + \left(\frac{4}{3} - \frac{4}{3k^2}\right)\beta^3 + \mathcal{O}\left(\beta^4\right) \end{aligned} \tag{S12}$$

The small-$\beta$ expansion of [1/2] Pade Approximant is

$$H(\beta) = 1 + \frac{A\beta}{1 + B\beta + C\beta^2} = 1 + A\beta - AB\beta^2 + A\left(B^2 - C\right)\beta^3 + \ldots$$

We will match the coefficients up to $\beta^3$ for the first 4 terms. We match the coefficients of $\beta, \beta^2, \beta^3$:

$$\begin{aligned} &A = h_1, \\ &-AB = h_2 \Longrightarrow B = -\frac{h_2}{A} = -\frac{h_2}{h_1}, \\ &A\left(B^2 - C\right) = h_3 \Longrightarrow C = B^2 - \frac{h_3}{A} = (\frac{h_2}{h_1})^2 - \frac{h_3}{h_1} \end{aligned}$$

So

$$\begin{aligned} H^{(1)}(\beta) &\approx 1 + \frac{h_1^{(1)}\beta}{1 - \frac{h_2^{(1)}}{h_1^{(1)}}\beta + [(\frac{h_2^{(1)}}{h_1^{(1)}})^2 - \frac{h_3^{(1)}}{h_1^{(1)}}]\beta^2} \\ H^{(2)}(\beta) &\approx 1 + \frac{h_1^{(2)}\beta}{1 - \frac{h_2^{(2)}}{h_1^{(2)}}\beta + [(\frac{h_2^{(2)}}{h_1^{(2)}})^2 - \frac{h_3^{(2)}}{h_1^{(2)}}]\beta^2} \end{aligned} \tag{S13}$$

where $h_1^{(1)} = \frac{2}{k}, h_2^{(1)} = -2 - \frac{2}{k} + \frac{2}{k^2}, h_3^{(1)} = -k^2 - \frac{8}{3}k - \frac{4}{3}$, and $h_1^{(2)} = 2, h_2^{(2)} = 2 - \frac{2}{k} - \frac{2}{k^2}, h_3^{(2)} = \frac{4}{3} - \frac{4}{3k^2}$. Below is an example comparing the exact simulation result at $k = -2.4$ from the main text, and our approximations (see figure S1, top panel). If we relax the condition and don't require that $\beta \to \infty, H(\beta) \to 1$. We can still use a [1/2] Padé approximant, but in the traditional form $H(\beta) = \frac{A\beta}{1+B\beta+C\beta^2}$. Diversity values below 1 are invalid and thus cut off(See Figure S1, bottom panel). We can also tune different orders of Padé approximants for higher accuracies.

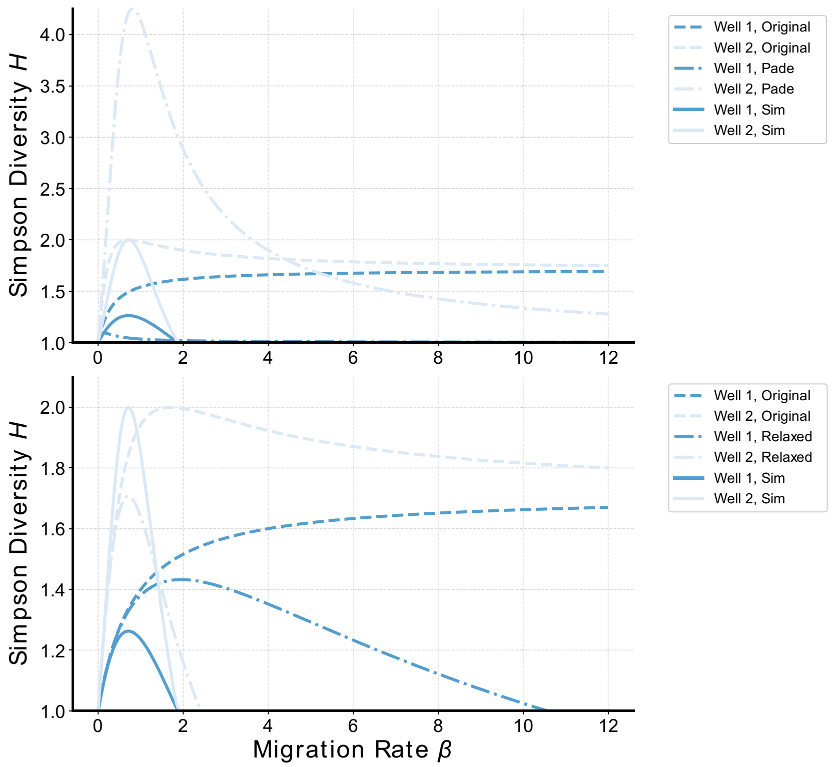


Figure S1: **An example comparing the exact simulation result at $k = -2.4$ from main text, and our approximations.** The top panel shows the comparison between exact simulation, the original approximation by small migration expansion, and the Padé approximation considering all restrictions. The bottom panel shows the relaxed Padé approximation calculated by python package *mpmath*.

##### Asymmetry parameter $k$ in the multiple-phenotype system

Here we show how asymmetry parameter $k = -\frac{g_{i*}^{(1)} - g_{j*}^{(1)}}{g_{i*}^{(2)} - g_{j*}^{(2)}} = \frac{\delta g_{i*}^{(1)}}{\delta g_{j*}^{(2)}}$ naturally emerges from the multi-phenotype system, and why it's still a proper shape parameter for diversity-migration relationship in 2 wells. For each phenotype $i$

in Well 1 and Well 2, the proportion dynamics are:

$$\frac{dp_i^{(1)}}{dt} = p_i^{(1)} \left( g_i^{(1)} - \sum_{j=1}^{N} x_j^{(1)} g_j^{(1)} \right) - \beta \left( p_i^{(1)} - p_i^{(2)} \right)$$

$$\frac{dp_i^{(2)}}{dt} = p_i^{(2)} \left( g_i^{(2)} - \sum_{j=1}^{N} p_j^{(2)} g_j^{(2)} \right) + \beta \left( p_i^{(1)} - p_i^{(2)} \right)$$

For N phenotypes, as $\beta = 0$, in each well it's still the phenotype with highest local fitness advantage that dominates. If each well has its own local dominant phenotype, we have $p_{i^*}^{(1)} = 1, p_{j^*}^{(2)} = 1, p_i^{(1)} = 0, p_j^{(2)} = 0, i \neq i^*, j \neq j^*, i^* \neq j^*$.

As $\beta$ increases but still small, we have (a) $p_{i^*}^{(1)} = 1 - \sum_{i \neq i^*} \delta p_i^{(1)}$, where $\delta p_i^{(1)} \ll 1$ represents the small fraction of non-dominant phenotypes in Well 1; (b) $p_{j^*}^{(2)} = 1 - \sum_{j \neq j^*} \delta p_j^{(2)}$, where $\delta p_j^{(2)} \ll 1$ represents the fraction of non-dominant phenotypes in Well 2; (c) $p_i^{(1)} = \delta p_i^{(1)}, p_j^{(2)} = \delta p_j^{(2)}, i \neq i^*, j \neq j^*$.

For local dominant phenotypes, let's start with phenotype $i^*$ in Well 1. knowing $p_{i^*}^{(1)}$ is close to 1. The growth-migration equation for phenotype $i^*$ in Well 1 is:

$$\frac{dp_{i^*}^{(1)}}{dt} = p_{i^*}^{(1)} \left( g_{i^*}^{(1)} - \sum_{j=1}^{N} p_j^{(1)} g_j^{(1)} \right) - \beta \left( p_{i^*}^{(1)} - p_{i^*}^{(2)} \right)$$

Rewrite the sum as $\sum_{j=1}^{N} p_j^{(1)} g_j^{(1)} = p_{i^*}^{(1)} g_{i^*}^{(1)} + \sum_{i \neq i^*} p_i^{(1)} g_i^{(1)}$. Substituting $p_{i^*}^{(1)} = 1 - \sum_{i \neq i^*} \delta p_i^{(1)}$ and $p_i^{(1)} = \delta p_i^{(1)}$, we get:

$$\sum_{j=1}^{N} p_j^{(1)} g_j^{(1)} = \left( 1 - \sum_{i \neq i^*} \delta p_i^{(1)} \right) g_{i^*}^{(1)} + \sum_{i \neq i^*} \delta p_i^{(1)} g_i^{(1)}$$

Substitute this back to the growth term and factor out $\sum_{i \neq i^*} \delta p_i^{(1)}$, it gives

$$p_{i^*}^{(1)} \sum_{i \neq i^*} \delta p_i^{(1)} \left( g_{i^*}^{(1)} - g_i^{(1)} \right)$$

Define $\Delta g_i^{(1)} = g_{i^*}^{(1)} - g_i^{(1)}$, the difference between the growth rates of the local dominant phenotype $i^*$ and the non-dominant phenotype $i$. Thus we have

$0 = p_{i^*}^{(1)} \sum_{i \neq i^*} \delta p_i^{(1)} \Delta g_i^{(1)} - \beta \left( \delta p_{i^*}^{(1)} - \delta p_{i^*}^{(2)} \right)$. Substitute $p_{i^*}^{(1)} = 1 - \sum_{i \neq i^*} \delta p_i^{(1)}$ again, ignore the higher-order terms like $(\delta p)^2$,

$$0 = \sum_{i \neq i^*} \delta p_i^{(1)} \Delta g_i^{(1)} - \beta \left( 1 - \sum_{i \neq i^*} \delta p_i^{(1)} - \delta p_{i^*}^{(2)} \right)$$

Do the re-arrangement and separate $p_{j^*}^{(1)}$. Finally we get

$$\sum_{i\neq i^*,j^*} \delta p_i^{(1)}(\Delta g_i^{(1)}+\beta) + \delta p_{j^*}^{(1)}(\Delta g_{j^*}^{(1)}+\beta) = \beta(1-\delta p_{i^*}^{(2)}) \tag{S14}$$

Similarly, for phenotype $j^*$ in Well 2,

$$\sum_{i\neq i^*,j^*} \delta p_i^{(2)}(\Delta g_i^{(2)}+\beta) + \delta p_{i^*}^{(2)}(\Delta g_{i^*}^{(2)}+\beta) = \beta(1-\delta p_{j^*}^{(1)}). \tag{S15}$$

For Non-dominant phenotypes, still, let's consider phenotype $i$ in Well 1 first. Let the left side of the selection-migration equation equal to 0 and substitute $p_i^{(1)} = \delta p_i^{(1)}, p_i^{(2)} = \delta p_i^{(2)}, i \neq i^* \neq j^*$, we have

$$\begin{aligned}
0 &= \delta p_i^{(1)} \left( g_i^{(1)} - \sum_i \delta p_i^{(1)} g_i^{(1)} \right) - \beta \left( \delta p_i^{(1)} - \delta p_i^{(2)} \right) \\
&= \delta p_i^{(1)} \left( g_i^{(1)} - \sum_{i\neq i^*} \delta p_i^{(1)} g_i^{(1)} - (1 - \sum_{i\neq i^*} \delta p_i^{(1)}) g_{i=i^*}^{(1)} \right) - \beta \left( \delta p_i^{(1)} - \delta p_i^{(2)} \right) \\
&\approx -\delta p_i^{(1)} \Delta g_i^{(1)} - \beta \left( \delta p_i^{(1)} - \delta p_i^{(2)} \right)
\end{aligned}$$

The last equality is by ignoring higher-order terms $(\delta p)^2$. Do the re-arrangement and we get

$$\delta p_i^{(1)} = \frac{\beta \delta p_i^{(2)}}{\Delta g_i^{(1)} + \beta} \tag{S16}$$

Similarly for non-dominant phenotype in Well 2,

$$\delta p_i^{(2)} = \frac{\beta \delta p_i^{(1)}}{\Delta g_i^{(2)} + \beta} \tag{S17}$$

By observation, $\delta p_i^{(1)} = \delta p_i^{(2)} = 0$ or $\frac{\beta}{\Delta g_i^{(1)}+\beta}\frac{\beta}{\Delta g_i^{(2)}+\beta} = 1, \beta = -\frac{\Delta g_i^{(1)}\Delta g_i^{(2)}}{\Delta g_i^{(1)}+\Delta g_i^{(2)}} < 0$. So under small $\beta$ limit, all non-dominant phenotypes in both wells are approximately 0. Thus the equations for dominant phenotypes become

$$\begin{aligned}
\delta p_{j^*}^{(1)}(\Delta g_{j^*}^{(1)} + \beta) &\approx \beta(1-\delta p_{i^*}^{(2)}) \\
\delta p_{i^*}^{(2)}(\Delta g_{i^*}^{(2)} + \beta) &\approx \beta(1-\delta p_{j^*}^{(1)})
\end{aligned} \tag{S18}$$

Rewriting $p_1 = 1-\delta p_{j^*}^{(1)}, p_2 = \delta p_{i^*}^{(2)}$, it's effectively a 2-phenotype system as discussed. We can thus define asymmetry parameter $k = \frac{g_{i^*}^{(1)} - g_{j^*}^{(1)}}{g_{i^*}^{(2)} - g_{j^*}^{(2)}} = \frac{\Delta g_{i^*}^{(1)}}{\Delta g_{j^*}^{(2)}}$(or replace $\Delta$ by $\delta$ to make it consistent with the symbol in main text). Although multiple species are competing with each other in both wells, we can still use a single asymmetry parameter $k$ to captures the main feature, determined only by the growth information from 2 local dominant phenotypes. Letting $\Delta g_{j^*}^{(1)} = 1$ without loss of generality, it returns back to the 2-phenotype scenario where

$$\begin{aligned} \delta p^{(1)} &\approx \frac{\beta}{|k| + \beta(|k|+1)} \\ \delta p^{(2)} &\approx \frac{|k|\beta}{|k| + \beta(|k|+1)} \end{aligned} \tag{S19}$$

at $k < 0$. We can still get $H_1, H_2$ as in the 2-phenotype scenario, by ignoring the higher-order terms, and use the Pade approximation.

So as $\beta$ is still small enough, we can reduce N phenotypes to 2 effective phenotypes $i^*, j^*$, since only they have large enough fitness to support the invasion in other wells. Thus we can define the asymmetry parameter $k = \frac{g_{i^*}^{(1)} - g_{j^*}^{(1)}}{g_{i^*}^{(2)} - g_{j^*}^{(2)}}$. This is a first-order approximation and it still works as an indicator to tell us the shape of the diversity-migration relationship - if it's satisfying "Intermediate Disturbance Hypothesis", and which well will have larger diversities. However, it cannot predict the multimodalities at the intermediate migration level, which need considerations of higher-order asymmetry parameters, by the other phenotypes which behave like generalists.

##### Perfect permutation induces singular behavior starting from a 4-phenotype system

To give more illustrations on the emergence of singular behaviors like multiple peaks of the diversity-migration relationship from the multi-phenotype system, we define the "perfect permutation": given an ordered growth rate set for phenotypes in one well, if we can obtain another growth rate set for phenotypes in the second well simply by permuting the original set, we call it a "perfect permutation." For example, in a 2-phenotype system, if the growth rates in well 1 are $\{g_1, g_2\}$ for phenotypes 1 and 2, then the only permuted growth rates in well 2 would be $\{g_2, g_1\}$. This corresponds to the case where $k = -1$, and the diversity-migration relationship in the two wells will be the same (see Figure S2A, left panel). Similarly, for a 3-phenotype system, there are three different permutations. Here, we only consider the effective permutations and ignore cases equivalent to the 2-phenotype system where the smallest growth rate is not permuted. For example, $\{g_1, g_2, g_3\}$ and $\{g_1, g_3, g_2\}$ are ignored. We observe that there are two cases satisfying the intermediate disturbance hypothesis with maximum diversity at different wells, while the third case is similar to the results from the 2-phenotype system. Specifically, under this scenario, only the largest and smallest growth rates are permuted (see Figure S2A, right panel). Therefore, no singular behaviors are observed in the 3-phenotype system.

For 4-phenotype systems, we still consider the effective permutations, resulting in 24 different cases. Interestingly, the 13th and 20th cases exhibit multiple peaks, although with a similar large migration behavior to that observed in 2-phenotype systems (Figure S2B). These cases involve growth rates $\{g_2, g_4, g_3, g_1\}$ and $\{g_4, g_1, g_3, g_2\}$ in well 2, with $\{g_1, g_2, g_3, g_4\}$ in well 1. We can, of course, introduce small perturbations to the permuted growth rates, and in this scenario, the minimal example becomes a generalist-specialist framework with three phenotypes, as shown in the main text.

##### Largest eigenvalue and global fitness advantage

Instead of defining higher-order asymmetry parameters, here we show that how global fitness advantage predicts the singular behaviors like multimodalities at the intermediate migration level. Before that, let's first discuss the largest eigenvalue of the original growth-migration dynamics.

For our growth-migration dynamics, consider each phenotype $i$ in 2 well system has the following form

$$\begin{gathered}\frac{\partial u_i}{\partial t} = -\beta L \mathrm{u}_i + G_i u_i = \Omega_i u_i \\ i = 1, 2, \cdots N\end{gathered} \tag{S20}$$

where $\mathbf{\Omega_i} = \begin{pmatrix} g_i^{(1)} - \beta & \beta \\ \beta & g_i^{(2)} - \beta \end{pmatrix}$

Here we focus on one phenotype and omit the subscript $i$ for simplicity. This is a linear system and the solution is $u = u_0 e^{\mathbf{\Omega} t}$. We can diagonalize it and get

$$\mathbf{u}(t) = c_1 e^{\lambda_1 t}\mathbf{u}_1 + c_2 e^{\lambda_2 t}\mathbf{u}_2,$$

where $c_1 =< \mathbf{u_0}|\mathbf{u_1} >$ and $c_2 =< \mathbf{u_0}|\mathbf{u_2} >$. $\mathbf{u}_1, \mathbf{u}_2$ are eigenvectors of the corresponding eigenvalues $\lambda_1, \lambda_2$ of $\mathbf{\Omega}$. For a 2-dimensional system, the characteristic polynomial is of the form $\lambda^2 - \tau\lambda + \Delta = 0$ where $\tau$ is the trace and $\Delta$ is the determinant of $\mathbf{\Omega}$. Thus the two roots are in the form:

$$\lambda_1 = \frac{\tau + \sqrt{\tau^2 - 4\Delta}}{2}$$
$$\lambda_2 = \frac{\tau - \sqrt{\tau^2 - 4\Delta}}{2}$$

So the exact solutions of eigenvalues and eigenvectors are

$$\begin{aligned}\lambda_1 &= \frac{1}{2}(-2\beta + g^{(1)} + g^{(2)} + \sqrt{4\beta^2 + (g^{(1)})^2 - 2g^{(1)}g^{(2)} + (g^{(2)})^2}), \\ \lambda_2 &= \frac{1}{2}(-2\beta + g^{(1)} + g^{(2)} - \sqrt{4\beta^2 + (g^{(1)})^2 - 2g^{(1)}g^{(2)} + (g^{(2)})^2})\end{aligned} \tag{S21}$$

$$\begin{aligned}\mathbf{u_1} &= (-\frac{-g^{(1)} + g^{(2)} - \sqrt{4\beta^2 + (g^{(1)})^2 - 2g^{(1)}g^{(2)} + (g^{(2)})^2}}{2\beta}, 1)^T, \\ \mathbf{u_2} &= (-\frac{-g^{(1)} + g^{(2)} + \sqrt{4\beta^2 + (g^{(1)})^2 - 2g^{(1)}g^{(2)} + (g^{(2)})^2}}{2\beta}, 1)^T\end{aligned} \tag{S22}$$

We have the first component of $\mathbf{u_1}$ is positive and the first component of $\mathbf{u_2}$ is negative. We can also find out that, as $t \to \infty$, we have $\mathbf{u}(t) \approx c_1 e^{\lambda_1 t}\mathbf{u}_1$. We can also see that $c_1 > 0, c_2 < 0$. If $\beta > \beta_c = F(g^{(1)}, g^{(2)})/2$(where $F(g^{(1)}, g^{(2)}) = \frac{2}{\frac{1}{g^{(1)}}+\frac{1}{g^{(2)}}}$ is the harmonic mean), $\lambda_2$ becomes negative. This is the critical point where only one effective growth rate $\lambda_1$ contains the growth information across both wells. The growth rate for this specific phenotype starts becoming global. Interestingly for each phenotype $i$ it has its own critical point $\beta_{c,i} = \frac{F(g_i^{(1)}, g_i^{(2)})}{2}$. The system remains dominated by 2 local dominant phenotypes as long as $\beta$ does not exceed the minimum of all $\beta_{c,i}$ :

$$\beta_{\min} = \min \left\{ \frac{F(g_i^{(1)}, g_i^{(2)})}{2} \right\}_{i=1}^{N}$$

This value is usually small according to experimentally measured growth rates. We can also imagine that, as $\beta < \beta_{\min}$, we have our diversity approximation valid. This also tell us that, it's potential for the diversity-migration relationship to have up to $N-1$ Peaks: Each phenotype can introduce a peak corresponding to its critical migration rate. Although in nature, the number of peaks is often fewer due to fluctuation or overlapping critical migration rates and species interactions.

To show how global fitness advantage emerges from this $\lambda_1$, let's go back to its function form

$$\lambda_1 = \frac{1}{2}\left(-2\beta + g^{(1)} + g^{(2)} + \sqrt{4\beta^2 + (g^{(1)})^2 - 2g^{(1)}g^{(2)} + (g^{(2)})^2}\right),$$

An interesting case here is, if $g^{(1)} = g^{(2)} = g$ as spatial homogeneity, $\lambda_1 = \frac{g^{(1)}+g^{(2)}}{2} = g$ showing a very simple form. Doing an large-$\beta$ expansion of $\sqrt{4\beta^2 + \left(g^{(1)} - g^{(2)}\right)^2}$,

$$\begin{aligned}\lambda_1 &\approx \frac{1}{2}\left[g^{(1)} + g^{(2)} + \frac{\left(g^{(1)} - g^{(2)}\right)^2}{4\beta} - \frac{\left(g^{(1)} - g^{(2)}\right)^4}{64\beta^3} + \cdots\right] \\ &\approx \frac{g^{(1)} + g^{(2)}}{2} + \frac{\left(g^{(1)} - g^{(2)}\right)^2}{8\beta} + \mathcal{O}\left(\frac{1}{\beta^3}\right)\end{aligned} \tag{S23}$$

As $\beta \to \infty$, if we only keep the first term we have $\lambda \approx \frac{g^{(1)}+g^{(2)}}{2} = g^{global}$ and our global fitness advantage emerges from $\lambda_1$. And here we can construct the global fitness advantage as migration is really large, just by experimental measured growth rates. For the phenotype with global fitness advantage, it needs to satisfy $i^* = \underset{i}{\text{argmax}} g_i^{global}$. Thus $g^{global}$ can also be used as a parameter to tune bimodal peaks, since it can also define a transition of 2 different global dominant phenotypes, similar to $\beta_c$. And we can use only 3 phenotypes such as 1 generalist and 2 specialists as a minimal example shown in main text.

##### Equivalence between model of exponential growth, and model of competition

Non-interacting population we used in this paper, or exponential growth, is a simplified model; in the experiments, population cannot increase infinitely, and we have the constraint of intra-phenotype and inter-phenotype competition, as described in a logistic model. At carrying capacity, we can treat the fractions of different phenotypes as unchanged. However, we can also prove that, if the carrying capacity is ”positively growth-dependent” (which is common for some cases where drugs modulate carrying capacities[4]), then our exponential growth model is equivalent to the logistic growth model. Assume we have the microscopic dynamics describing logistic growth and migration and use 2 phenotype sensitive(S) and resistant(R) in well 1 as an example

$$\frac{du_S^{(1)}}{dt} = u_S^{(1)}(g_S^{(1)} - \frac{g_S^{(1)}u_S^{(1)} + g_R^{(1)}u_R^{(1)}}{K}) + \beta u_S^{(2)} - \beta u_S^{(1)} \tag{S24}$$

If we rescale both sides by carrying capacity $K$ and define $p = \frac{u}{K}$ as the fraction, then we have

$$\frac{dp_S^{(1)}}{dt} = p_S^{(1)}(g_S^{(1)} - (g_S^{(1)}p_S^{(1)} + g_R^{(1)}p_R^{(1)})) + \beta p_S^{(2)} - \beta p_S^{(1)} \tag{S25}$$

which is exactly

$$\frac{dp_S^{(1)}}{dt} = p_S^{(1)}(g_S^{(1)} - \langle g^{(1)}\rangle) + \beta p_S^{(2)} - \beta p_S^{(1)} \tag{S26}$$

the same form as the exponential growth in this paper. Since we know that dilution doesn't affect the relative abudance (fraction), our exponential growth model can well capture the experimental dynamics under this scenario.

#### Other diversity metric

For $q = 1$, ${}^qH = \left(\sum_{i=1}^S p_i^q\right)^{1/(1-q)}$ is undefined, but its limit as $q \to 1$ is the exponential of the familiar Shannon diversity:

$${}^1H = \lim_{q\to 1} {}^qH = \exp\left(-\sum_{i=1}^{S} p_i \log p_i\right)$$

The variable ${}^1H$ weighs species in proportion to their frequency[5, 6]. Shannon entropy is also widely used in describing diversity-disturbance relationship[7], and usually it shows minor differences comparing to Simpson diversity as we chose in our paper[8].

### Experiment-related methods and supplementary results

#### Experimental procedure for the migration-evolution lab system

Here we illustrate how our migration-evolution lab system generally works. 8 biological replicates of isogenic E. faecalis were grown overnight. 2 columns from a 96 deep well plate were chosen as 2 patches for 8 replicates. Different drug concentrations were added to different pairs of 2 columns at drug wells. After the bacteria were added, well plates were put into the incubator for a daily-based phenotype selection. Starting from the second day(or day 1 in our growth-migration-dilution cycle), a 8-channel pipetting robot performed the mixing between 2 wells, with different mixing times to mimic different migration rates. Then dilution would be performed with new well plates, to minimize the drug mixing effect in the migration step. The same daily cycle was repeated for 8 days. IC50s were later measured for resistant phenotype classification(see Figure S3).

#### Sanity check of AMP degradation over 8 days

In our migration-evolution experiments, we prepared ampicillin drug stocks before initiating the short-term evolution phase. While ampicillin is known to degrade over time, even when stored at -80°C, such degradation could potentially influence the outcomes of the evolutionary experiment. To ensure the reliability of our results, we conducted a sanity check to monitor ampicillin degradation as well as to assess potential fluctuations in the IC50 of wild-type (WT) strains. Specifically, we measured the AMP IC50 of eight WT biological replicates over eight consecutive days, using ampicillin stocks prepared on day 0.

The results show that there is no increasing trend in AMP IC50 values over time, indicating that ampicillin degradation is unlikely to have impacted the experimental conditions. Although some fluctuations were observed in the daily IC50 values, the average WT IC50 for AMP remained consistent across the experiment. Importantly, these values remained well within the expected range and did not approach the threshold lines representing twice or half of the average IC50 (see Figure S4). This suggests that the observed IC50 variations are due to normal biological variability rather than drug degradation.

#### Lack of cross resistance in drug-media experiments

In the AMP-LZD experiment, we identified the existence of a cross-resistance (RR) phenotype, which is resistant to both ampicillin and linezolid. Since it is unclear whether this phenotype results from collective drug selection or simply from the effects of either ampicillin or linezolid alone, we conducted supplementary AMP IC50 measurements on replicates from the LZD-media experiment.

Similarly, we performed LZD IC50 measurements on replicates from the AMP-media experiment. We assumed that there are no strong collateral effects, such as enhanced sensitivity or resistance in phenotypes selected by either AMP or LZD individually. According to recent research, for V583, a commonly used laboratory strain of E. faecalis, AMP-selected phenotypes exhibited slight collateral resistance to LZD, while LZD-selected phenotypes showed slight collateral sensitivity to AMP [9]. However, these effects were not pronounced.

Our results show that for strains selected in drug-media experiments, there is no evidence of significant cross-resistance (high IC50s for both drugs, which would be represented in the upper right corner of the left panel, Figure S5) or collateral sensitivity. The log2-log2 scatter plot is a common visualization for distinguishing different resistance levels in antibiotic resistance evolution[9]. While some replicates exhibit cross-resistance or collateral sensitivity, most values fall within a reasonable range, between half and twice the average WT IC50s (Figure S5, left panel). In the log-log scatter plot of strains from the AMP-LZD experiment, there is clear evidence of multiple replicates clustering around the high AMP, high LZD resistance region (Figure S6, left panel).

Comparing the histograms of LZD and AMP IC50s between drug-media and two-drug experiments (Figure S5 and Figure S6, right panels), we observe higher resistance levels to AMP and LZD in the two-drug experiments. This suggests that the presence of both drugs selects more effectively for cross-resistance or higher resistance levels to individual drugs. This effect can be explained by the absence of sanctuaries in the two-drug environment, where cross resistance phenotypes have more fitness advantages; also strains selected by both drugs tend to incur greater fitness costs, making them less optimal for survival under single-drug selection. Besides, we can conclude that the "time-delayed" responses from AMP-media experiments, is not due to an extra phenotype like cross resistance.

#### IC50 Distributions of Isolated Samples

In the two-drug experiments, although we identified cross-resistance, the IC50s were measured from population-level samples. To determine whether this cross-resistance was due to a mixed population of AMP- and LZD-resistant cells or represented true cross-resistance phenotypes, we also assessed the IC50 distributions at the individual-cell level. Additionally, we aimed to understand how well population-level IC50s align with individual-cell level IC50s and to quantify the extent of fluctuation.

To obtain individual-cell level IC50s, we regrew candidate strains on agar plates and selected 32 isolates for each strain. These isolates represent IC50 values from a microscopic perspective, as each isolate originates from a single cell that grew and expanded on the agar plate. We designated each strain with a short name based on migration rate, drug concentration conditions, and replicate number, such as "migration-selecting drug-replicate." For example, "beta10L5R8" represents a strain selected under migration rate $\beta = 10$, in the LZD well with 5/6 MIC as the selecting drug concentration, and is the 8th biological replicate.

We selected 52 representative strains, encompassing four different phenotypes. The results show that most of the IC50 distributions are concentrated, though a few exhibit distinct resistance levels(Figure S7). For example, strains like beta5A5R5, beta5L5R6, and beta10A6R2 show two distinct LZD resistance levels. This could be attributed to the common point mutations on the 23S rDNA of E. faecalis under LZD selection [9, 10, 11]. Interestingly, strain beta5L5R4 also exhibits two distinct AMP resistance levels.

For cross-resistance, strains such as beta5L4R6, beta5A4R7, beta5L4R7, beta5L5R1, and beta5A5R5 present a possible mix of lower LZD resistance and higher LZD resistance. However, most of these strains are also AMP-resistant, indicating that they are true cross-resistance phenotypes with two LZD resistance clusters. Although we could introduce a "higher" LZD-resistant phenotype into the model, our minimal model predictions capture the main features observed in the experiments (see main text), so we retained the four phenotypes in the model as an effective representation.

Overall, the results indicate a general match between individual- and population-level IC50s, though discrepancies exist in strains like beta10L4R1 and beta10L6R7. The IC50 distributions at the individual-cell level reveal the complex resistance dynamics in two-drug evolution. Further research, both theoretical and experimental, is needed for a deeper understanding of these resistance mechanisms.

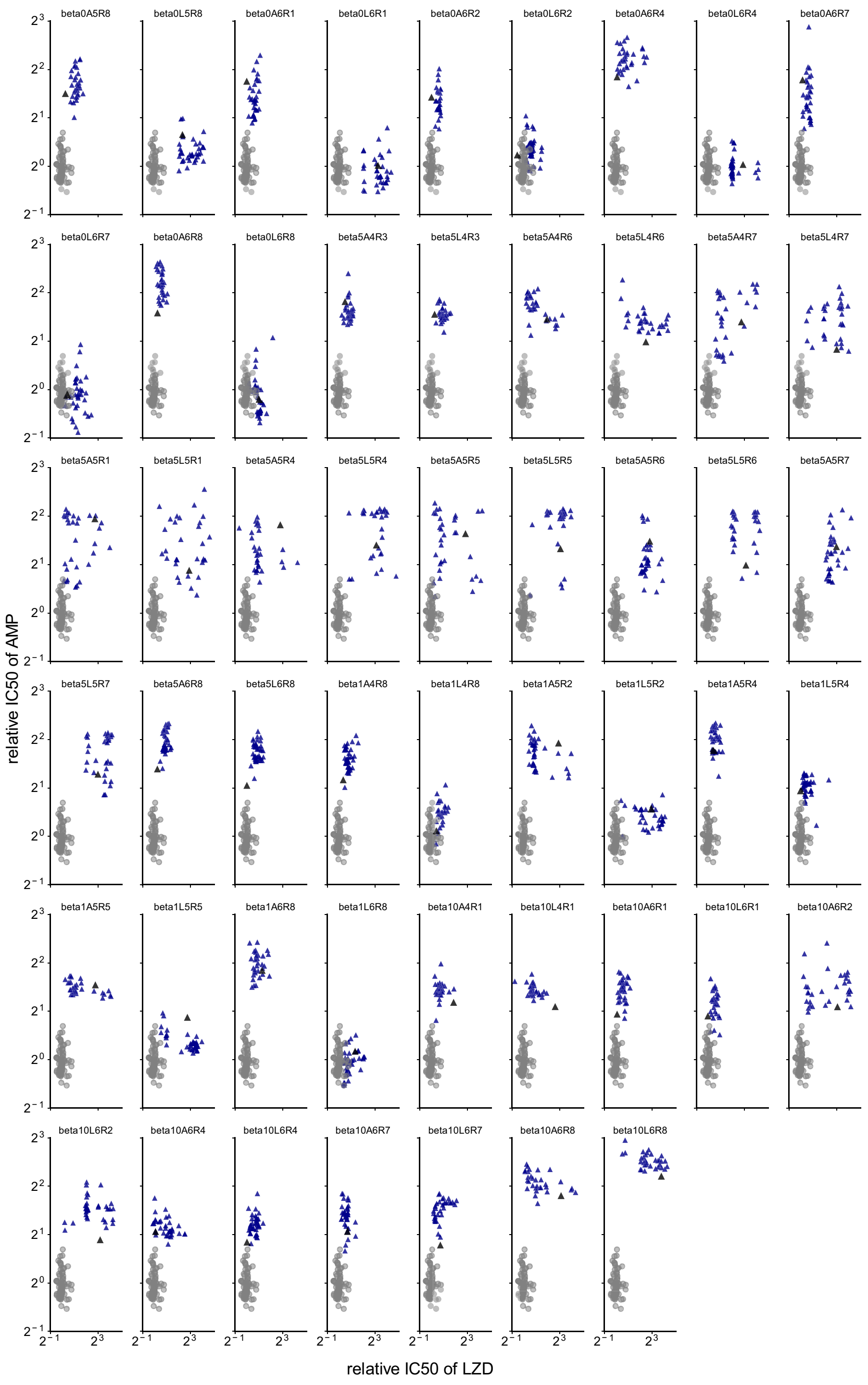
beta0A5R8
beta0L5R8
beta0A6R1
beta0L6R1
beta0A6R2
beta0L6R2
beta0A6R4
beta0L6R4
beta0A6R7
beta0L6R7
beta0A6R8
beta0L6R8
beta5A4R3
beta5L4R3
beta5A4R6
beta5L4R6
beta5A4R7
beta5L4R7
beta5A5R1
beta5L5R1
beta5A5R4
beta5L5R4
beta5A5R5
beta5L5R5
beta5A5R6
beta5L5R6
beta5A5R7
beta5L5R7
beta5A6R8
beta5L6R8
beta1A4R8
beta1L4R8
beta1A5R2
beta1L5R2
beta1A5R4
beta1L5R4
beta1A5R5
beta1L5R5
beta1A6R8
beta1L6R8
beta10A4R1
beta10L4R1
beta10A6R1
beta10L6R1
beta10A6R2
beta10L6R2
beta10A6R4
beta10L6R4
beta10A6R7
beta10L6R7
beta10A6R8
beta10L6R8
relative IC50 of AMP
relative IC50 of LZD

Figure S7: **Relative IC50 distributions of 32 isolates from 52 different strains on a log2 scale.** Grey dots represent the IC50 values of wild-type (WT) isolates. Dark blue triangles indicate IC50 values of individual isolates from the selected strain, while the black triangle represents the population-level IC50 for that strain.

#### Tolerance test for AMP: population-Level and single-Isolate analysis

To identify possible tolerance, we performed a growth-death experiment for all strains from the AMP-media experiment [12]. Strains exhibiting tolerance are expected to show no difference in AMP IC50 compared to the wild-type (WT), but would have lower drug-free growth rates and lower maximum lysis rates [13]. Our results indicate that almost all strains had lower growth rates and lysis rates compared to WT (Figure S8, left panel). Although AMP resistance can also lead to reduced drug-free growth rates and lysis rates due to fitness costs, the presence of non-AMP-resistant strains among those tested suggests that some strains are indeed exhibiting tolerance. Therefore, the discrepancies between the theoretical and experimental results discussed in the main text may be due, at least in part, to the presence of tolerance strains.

We also sought to determine if tolerance strains were present in the AMP-LZD experiments and, more specifically, whether tolerance occurred at the individual-cell level. We randomly selected ten strains (beta0A5R8, beta0L5R8, beta1A4R8, beta1L4R8, beta1A5R4, beta1L5R4, beta1A6R8, beta1L6R8, beta1A5R5, beta1L5R5, beta1A5R2, beta1L5R2) for further analysis. The results were consistent with our earlier observations, showing lower growth and lysis rates compared to WT (Figure S8, right panel). This suggests that ampicillin tolerance is common in our migration-evolution lab system under various migration rates and drug concentration conditions.

Further research is needed, including new experimental designs and novel modeling approaches, to clarify the role of tolerance in evolution under spatial drug heterogeneity.

#### More experimental results in the AMP-LZD migration-evolution experiment

Dose response curves of four emerging phenotypes in the AMP-LZD migration-evolution experiment are presented for both AMP and LZD. For the AMP dose response curves, the RL phenotype shows the lowest growth rate, indicating slight collateral sensitivity compared to the sensitive phenotype(Figure S9, left panel). Although all three resistant phenotypes exhibit a fitness cost in the absence of AMP, their growth rates are still comparable to the WT. At high AMP concentrations, the RR and RA phenotypes achieve the highest growth rates.

For the LZD dose response curves, the RA phenotype demonstrates almost no fitness cost and behaves similarly to the sensitive phenotype(Figure S9, right panel). After an initial small fitness cost, the RL and RR phenotypes display high growth rates at high LZD concentrations. These diverse responses suggest that applying different drugs in separate wells could yield even more interesting results, warranting further research.

For the two-drug experiment, we also measured IC50 values for strains from day 1 and day 4 to observe the emergence of the four phenotypes and the temporal change in the diversity-migration relationship. The results indicate that at day 1, the diversity values for all six different drug concentrations are 1, showing no evidence of resistant phenotypes(Figure S10, top panel). By day 4, resistant phenotypes begin to emerge, leading to an increase in diversity, particularly in the LZD well at high migration rates(Figure S10, bottom panel). All other drug concentrations continued to show no change.

Combining these observations with the day 8 results from the main text, we conclude that there is a bottleneck phase for resistance selection during the first few days, followed by a rapid emergence of resistance. It may also be valuable to validate the temporal dynamics of diversity in the future through additional experimental studies.

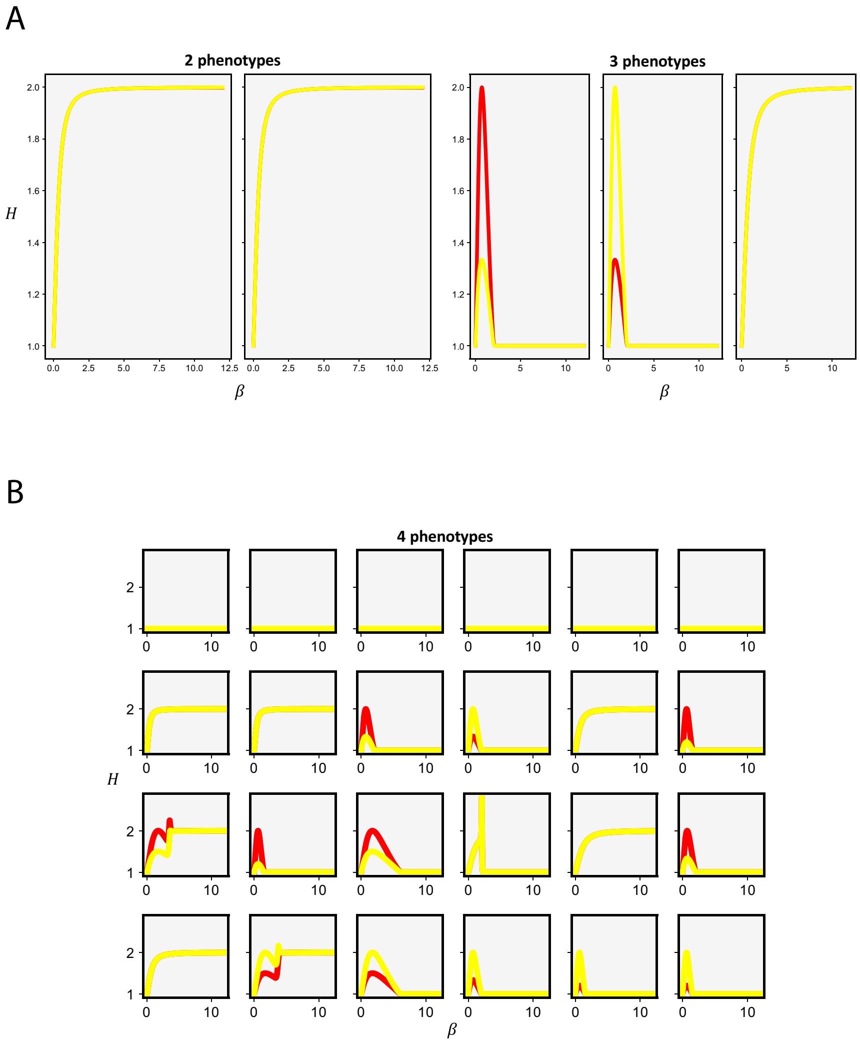


Figure S2: **Diversity-migration relationships by different perfect permutations, for 2,3,4 phenotypes.** Here we choose growth rate values $g = \{3.0, 4.0\}$ for 2 phenotypes, $g = \{2.0, 3.0, 4.0\}$ for 3 phenotypes, and $g = \{1.0, 2.0, 3.0, 4.0\}$ for 4 phenotypes.

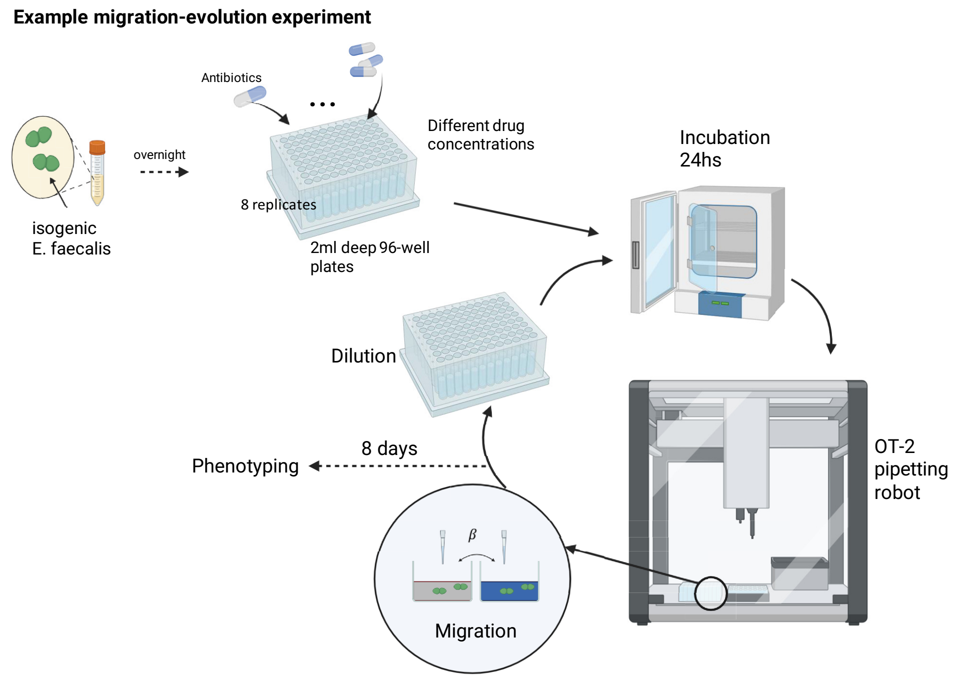


Figure S3: **Experimental procedure for the lab migration-evolution system.** A growth-migration-dilution cycle is repeated for each day until 8th day.

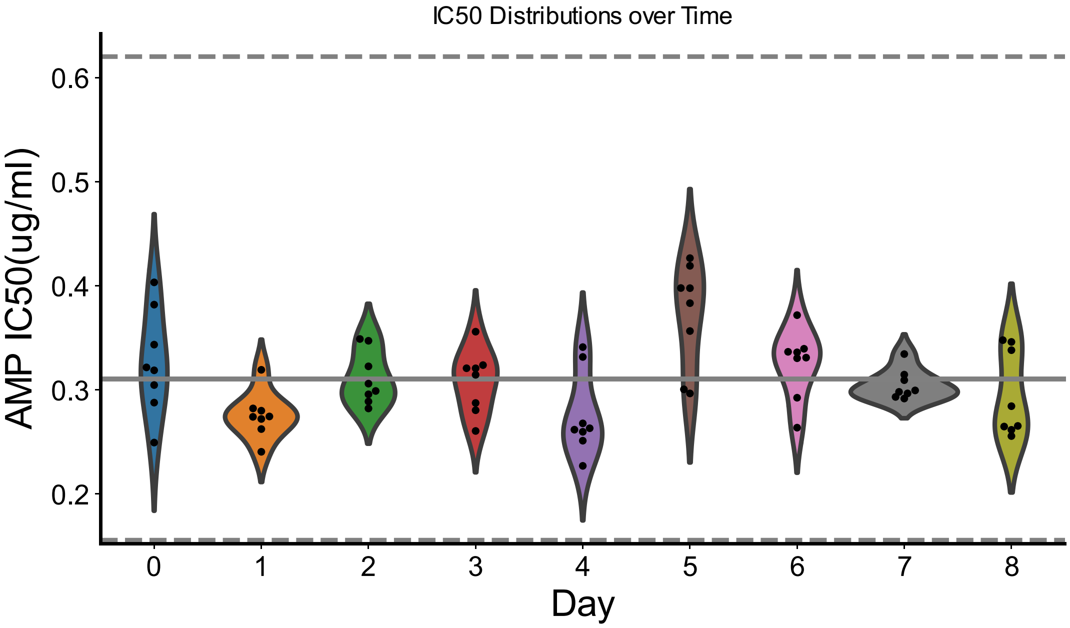


Figure S4: **AMP IC50s of WT at different days.** Dots represents different WT biological replicates. The solid line is the averaged AMP IC50. The top dashed line is twice of the average IC50 and the bottom dashed line is half the average IC50.

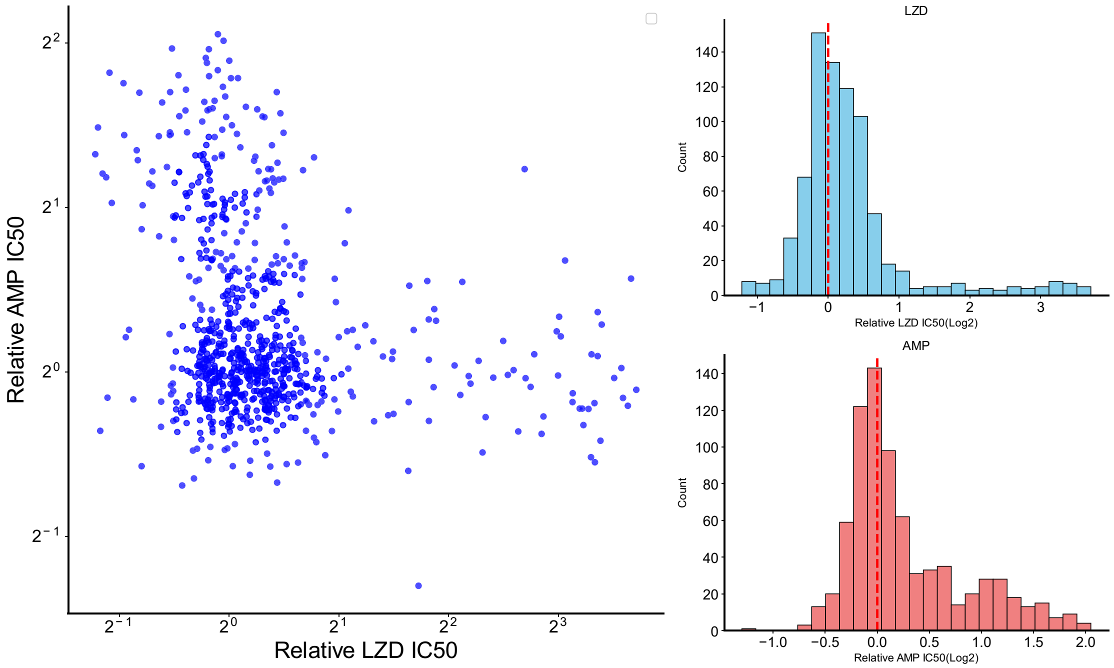


Figure S5: **Relative IC50s of AMP and LZD compared to WT for strains from drug-media experiments.** The log2-log2 scatter plot includes data from both AMP-media and LZD-media experiments. The column-like distribution of data points predominantly represents the AMP-media experiment, while the row-like distribution of data points mainly represents the LZD-media experiment. The two histograms on the right panel illustrate the distributions of AMP and LZD IC50s.

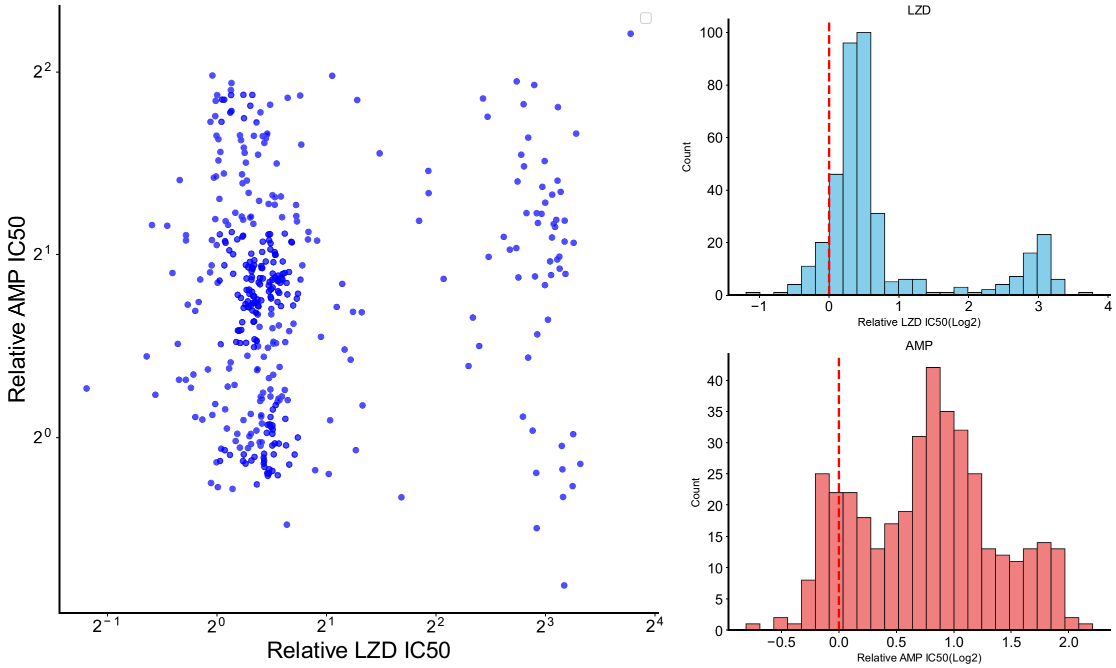


Figure S6: **Relative IC50s of AMP and LZD compared to WT for strains from AMP-LZD experiments.** The log2-log2 scatter plot shows evidence of cross-resistance phenotypes. The two histograms on the right panel illustrate the distributions of AMP and LZD IC50s.

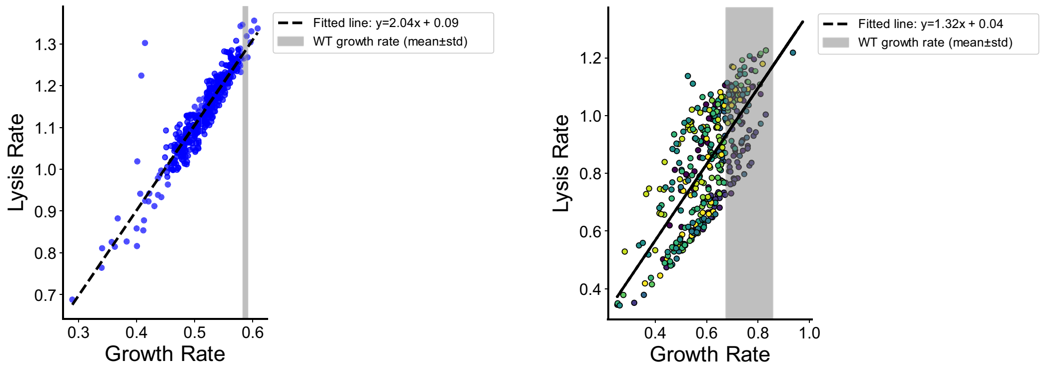


Figure S8: **Drug-free growth rates and maximum lysis rates for population-level strains from AMP-media experiments and individual-cell level strains from AMP-LZD experiments.** Data points in the left panel represent results from the AMP-media experiments, while data points in the right panel correspond to the AMP-LZD experiments. Both sets of data exhibit a strong linear correlation, and the slopes provide insights into how lysis rates can be inferred from growth rates. The grey shades represent normal ranges of WT growth rates and they are both on the right side.

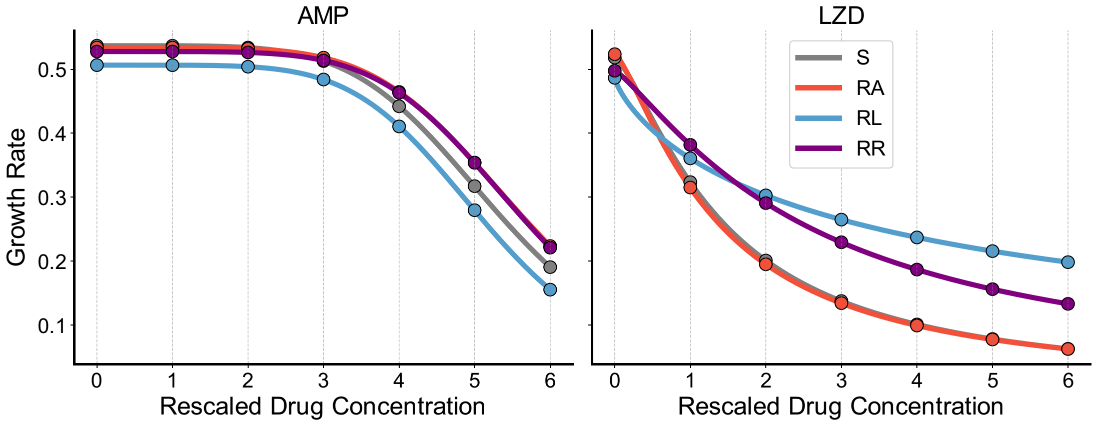


Figure S9: **Dose response curves of four emerging phenotypes from AMP-LZD experiments, for AMP and LZD.** Curves are fitted using all available growth rate data for the strains. A rescaled drug concentration of 6 represents the MIC.

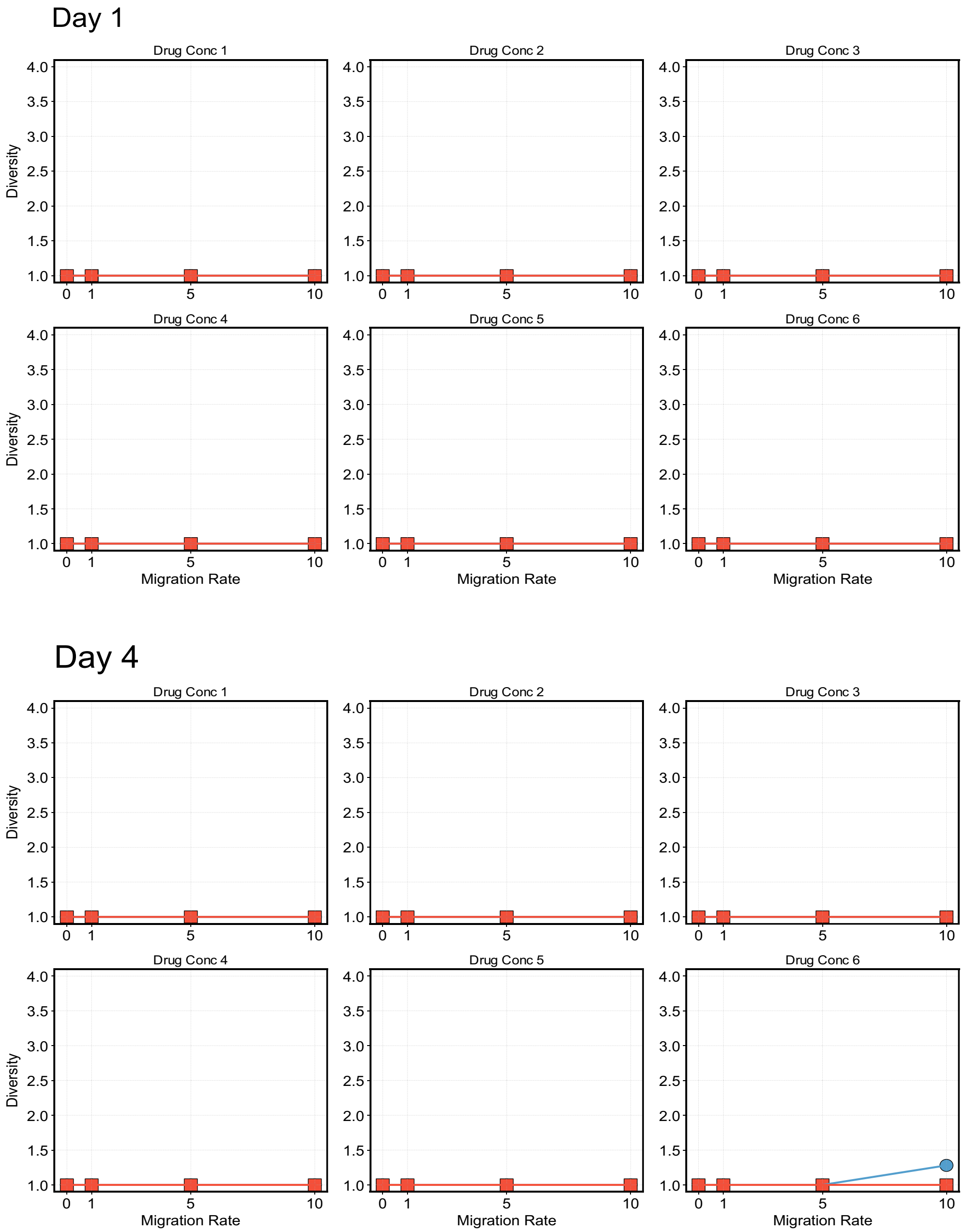


Figure S10: **Diversity-migration relationships for six different drug concentrations at day 1 and day 4.** Light red curves and dots represent AMP wells, while light blue curves and dots represent LZD wells.
